# Reward and punishment contingency shifting reveals distinct roles for VTA dopamine and GABA neurons in behavioral flexibility

**DOI:** 10.1101/2024.10.07.617060

**Authors:** Merridee J Lefner, Bita Moghaddam

## Abstract

In dynamic environments where stimuli predicting rewarding or aversive outcomes unexpectedly change, it is critical to flexibly update behavior while preserving recollection of previous associations. Dopamine and GABA neurons in the ventral tegmental area (VTA) are implicated in reward and punishment learning, yet little is known about how each population adapts when the predicted outcome valence changes. We measured VTA dopamine and GABA population activity while male and female rats learned to associate three discrete auditory cues to three distinct outcomes: reward, punishment, or no outcome within the same session. After learning, the reward and punishment cue-outcome contingencies were reversed, and subsequently rereversed. As expected, the dopamine population rapidly adapted to learning and contingency reversals by increasing the response to appetitive stimuli and decreasing the response to aversive stimuli. In contrast, the GABA population increased activity to all sensory events regardless of valence, including the neutral cue. Reversing learned contingencies selectively influenced GABA responses to the reward-predictive cue, prolonging increased activity within and across sessions. The observed valence-specific dissociations in the directionality and temporal progression of VTA dopamine and GABA calcium activity indicates that these populations are independently recruited and serve distinct roles during appetitive and aversive associative learning and contingency reversal.

## Introduction

In order to adaptively traverse an environment, one must learn to discriminate between cues that predict outcomes of different valences. The ventral tegmental area (VTA) is a central hub for acquiring associations as it contains a heterogeneous assembly of cells that respond to environmental stimuli, predominantly characterized as either dopamine or γ-aminobutyric acid (GABA) neurons^1-4^. Extensive research has established a fundamental role for VTA dopamine activity in processing rewarding stimuli^5-15^, although studies have determined that VTA dopamine neurons also exhibit responses to aversive stimuli^8,16-18^. Historically, investigation of VTA GABA activity has primarily focused on local modulation of dopamine function^19-24^. However, GABA neurons in the VTA also exert effects independent of VTA dopamine neurons^25-28^, and respond to appetitive and aversive stimuli in a manner that is distinct from dopaminergic activity^8,29,30^. This suggests that VTA dopamine and GABA populations are independently recruited during appetitive and aversive situations.

The ability to flexibly update behavior is essential for navigating dynamic environments where cues predicting rewarding or aversive consequences can unexpectedly change. To examine the role of VTA neurons in updating learned associations, our earlier work developed a flexible contingency learning (FCL) paradigm to assess the initial acquisition and subsequent reversal of positive and negative associations experienced within the same session^31,32^. Single unit recordings determined that VTA neurons were predominantly excited by appetitive predictive cues and inhibited by aversive predictive cues, with these signals dynamically updating following contingency reversal^31^. Correlated activity between VTA neurons increased to the appetitive association but decreased in response to the aversive association^32^, indicating that VTA dopamine and non-dopamine neurons produce separable responses throughout the reversal of contingencies. The response pattern of the VTA GABA population, as well as temporal differences between dopamine and GABA signaling across different phases of learning and reversal, however, remain poorly understood.

Here, we combined the FCL behavioral paradigm with fiber photometry recordings in male and female rats to track calcium influx dynamics in VTA dopamine and GABA populations throughout initial learning, reversal, and re-reversal of appetitive and aversive associations. We identified dissociable roles between VTA dopamine and GABA responses across FCL: increases in dopamine population calcium activity occurred in response to appetitive association, whereas the GABA population calcium activity increased to all stimuli. Additionally, reversing the contingences evoked a change in how GABA, but not dopamine, processed the new appetitive contingency. The GABA population activity increased towards the appetitive cue across reversal sessions, and within-session increases in GABA activity were dynamic during both early and late sessions of the reversal phase. These data collectively illustrate directional and temporal dissociations in VTA dopamine and GABA calcium activity to flexible reward and punishment contingencies.

## Methods

### Subjects

All procedures were approved by the Oregon Health & Science University Institutional Animal Care and Use Committee and were conducted in accordance with the National Institute of Health Guide for the Care and Use of Laboratory Animals. Male and female Long-Evans rats were bred in house on a TH::Cre (*n* = 8; 5 female) or GAD::Cre (*n* = 9; 5 female) transgenic background. All rats were given *ad libitum* access to water and chow, group-housed with littermates until aged to P60-65 when surgical procedures began, and maintained on a 12 h reverse light/dark cycle (lights off at 9:00 am) where all experiments were performed during the dark phase.

### Surgery

*Viral infusion surgery*. Rats were placed under isoflurane anesthesia and received unilateral or bilateral infusions of AAV1-Syn-Flex-GCaMP6f-WPRE-SV40 (1×10^13^vg/mL, Addgene) to allow for Cre-dependent expression of GCaMP in the VTA (AP -5.5 mm, ML +/-0.6 mm from bregma, DV -7.5, -6.5 mm from dura). Each infusion was 250 nL, administered at a rate of 100 nL/min, with the most ventral infusion performed first. The syringe was left in place for 10 min to allow virus to diffuse before slowly removing the needle. Animals were given 5 mg/kg of carprofen after surgery, after which they were single-housed.

*Fiber implant surgery*. After a minimum of 3 weeks following viral infusion surgery, subjects were implanted with a 400 μm diameter optical fiber targeting the VTA (AP -5.5 mm, ML +/-0.6 mm from bregma with left/right sides counterbalanced across animals, DV -7.3 mm from dura) along with 4 skull screws. Fibers were secured to the skull using a light-curing dental cement (Ivoclar Vivadent) followed by powder acrylic cement (Lang Dental). Subjects were administered 5 mg/kg of carprofen after surgery and were given at least 1 week to recover before behavioral testing began.

### Behavioral testing

After recovering from surgery, rats were placed and maintained on mild food restriction to target 90% free-feeding weight. Behavioral sessions were performed in conditioning chambers that included grid floors connected to a shock generator, a food trough, and three auditory stimulus generators (4.5 kHz tone, white noise, and clicker; Coulbourn Instruments). The chamber floors were thoroughly cleansed with disinfectant, and the walls and food port were cleaned with 70% ethanol solution between every subject. To familiarize rats with the chamber and food retrieval, rats underwent a single magazine training session in which 25 sucrose pellets (45 mg; BioServ) were noncontingently delivered at a 90 ± 15 s variable interval.

*Flexible contingency learning (FCL) task*. Rats were trained on a modified version of our previously developed FCL paradigm ^31,32^. In this task, subjects underwent 24 Pavlovian conditioning sessions in which the termination of a 5 s auditory cue [conditioned stimulus (CS); tone, white noise, or click] resulted in the delivery of an appetitive (sucrose pellet), aversive (0.2 mA, 180 ms shock), or neutral (nothing) outcome [unconditioned stimulus (US)]. All associations were presented in every session, and the CS-US pairings were counterbalanced across subjects. Each session contained 25 appetitive trials, 25 aversive trials, and 25 neutral trials delivered in a pseudorandom order, with a 45 ± 5 s intertrial interval between trials. After 8 sessions of initial training, the appetitive and aversive associations reversed in that the cue previously associated with the sucrose pellet (CS_1_) instead preceded the foot shock, and the cue previously associated with the foot shock (CS_2_) instead preceded the sucrose pellet. After 8 sessions of this reversal, rats underwent re-reversal where the appetitive and aversive cue-outcome associations returned to the original assignments (as experienced during initial training) for 8 final sessions. The neutral cue (CS-), which was associated with no outcome, did not change across sessions. Conditioned responding was quantified as the change in the rate of head entries to the food port during the 5 s CS relative to the 5 s preceding the CS delivery ^33^. We also quantified the latency to initiate a head entry during the CS, and the probability of initiating a head entry during the CS. For the post-outcome analysis, we calculated the average number of head entries made during a 5 s post-US delivery time window.

### Fiber photometry

Fiber photometry recordings for the detection of VTA dopamine or GABA population activity were performed in all sessions using a system with optical components from Doric lenses, with LED modulation controlled by a real-time processor from Tucker Davis Technologies (TDT; RZ5P). Rats were attached to a fiber optic cable in which 465 nm (signal) and 405 nm (isosbestic control) LEDs were modulated at 211 and 330 Hz, respectively, to the implanted cannula. The LED power was set for each animal to yield between 150–200 mV for each signal ^34^. Data was acquired with TDT Synapse software, and time stamps for the CS and US were collected via 5V TTL signals from the behavioral chamber that interfaced with the TDT processor in order to align events with calcium activity.

Analyses of GCaMP signals were performed using custom Python scripts based on those previously described ^35-37^. The isosbestic control and signal channels were low pass filtered at 3 Hz using a butterworth filter to reduce noise, and then the isosbestic control channel was fitted to the signal channel using a least squares polynomial fit of degree 1. Data was then separated into epochs based on the start and end of a given trial. The change in fluorescence (ΔF/F) was calculated by subtracting the fitted isosbestic control channel from the signal channel before dividing by the fitted isosbestic control channel. The signal was then z-scored by subtracting the mean ΔF/F from the ΔF/F signal, divided by the standard deviation of the ΔF/F signal. The z-score comparison window was the 5 s prior to the CS onset. To quantify signal changes in response to each CS, the average z-score was calculated during the entire 5 s CS, as well as during the first 2 s and last 3 s of the CS. To quantify responses to the US, the average z-score was calculated during the 3 s following US delivery, and the peak US response was calculated by taking the maximum z-score during the 3 s following CS termination relative to 0.5 s before US delivery. We additionally analyzed data in 5-trial bins in order to assess changes in calcium activity between the beginning and end of a session. To determine whether within-session changes in calcium activity occurred during early or late training sessions, the resulting binned data was averaged across the first 4 or last 4 sessions of each training phase.

### Fiber photometry permutation analysis

We used a permutation-based approach to compare changes in neural calcium activity as described previously^35,38^ using Python. For each subject and session, a z-score response to the appetitive, aversive, and neutral associations were separately calculated. For each comparison, a null distribution was generated by shuffling the data, randomly selecting the data into two groups, and calculating the mean difference between groups. This was performed 1000 times for each time point. A p-value was obtained by determining the percentage of times a value in the null distribution was greater than or equal to the observed difference in the unshuffled data (two-tailed for all comparisons). To control for multiple comparisons we utilized a consecutive threshold approach based on the 3 Hz lowpass filter window^35,38,39^, where a p-value < 0.05 was required for 14 consecutive samples in order to be considered significant.

### Data analysis and statistics

Detailed results of all statistical tests are found in the **Statistical Tables**. Aside from permutation tests, all statistical analyses were performed in GraphPad Prism 10. FCL behavioral responding, quantification of neural data, and binned trial data were analyzed using a 2-way mixed-effects model fit (restricted maximum likelihood method), repeated measures where appropriate, followed by *post hoc* Tukey’s or Sidak’s tests. The Geisser– Greenhouse correction was applied to address unequal variances between groups. Unpaired t-tests were used to compare open field data between groups.

### Histology

After behavioral testing, rats were deeply anesthetized with chloral hydrate (400 mg/kg, i.p.), then transcardially perfused with phosphate-buffered saline followed by 4% paraformaldehyde. Brains were removed and postfixed for >24 h, then subsequently placed in 30% sucrose solution. Sections were cut at 40 microns on a cryostat (Leica Microsystems) and stored in phosphate-buffered saline (PBS) with 0.05% sodium azide. Immunohistochemistry was performed to verify localization of GCaMP6f viral expression in VTA dopamine or GABA neurons for fiber photometry experiments, or to verify GiDREADD/control mCherry and CAV-cre expression for chemogenetic experiments. Brain slices were first permeabilized in 3% bovine serum albumin (BSA), 0.1% Triton X, and 1% Tween 80 in PBS + 0.05% sodium azide for 2 h at room temperature.

Sections were then incubated with the primary antibody mouse anti-GFP (1:500, Abcam) as well as either rabbit anti-glutamate decarboxylase (GAD; 1:500, Abcam) or chicken anti-tyrosine hydroxylase (TH; 1:500, Abcam) for fiber photometry experiments, diluted in PBS + Azide, 3% BSA + 0.15% Triton X for 24 h at 4°C. Slices were then washed in PBS + azide, 3% BSA + 0.15% Triton X, three times for five minutes each. Brain sections were next incubated with the secondary antibodies donkey-anti-mouse Alexa-488 (1:1000, Abcam), goat-anti-chicken Alexa-594 (1:1000, Abcam), and goat-anti-rabbit Alexa-594 (1:1000, Abcam), diluted in PBS + Azide, 3% BSA + 0.15% Triton X for 2 h at room temperature. Sections were washed again as outlined above and mounted to slides with Vectashield anti-fade mounting medium (Vector Labs). Brain slices were imaged for viral expression and fiber placement on a Zeiss Axio Observer microscope.

## Results

### Behavioral responding differentiates between associative valence and adapts to contingency reversal

Rats were trained on a flexible contingency learning (FCL) paradigm that was modified from our previous work^31,32^ to allow for the simultaneous acquisition of three distinct conditioned associations. In each session, 3 discrete auditory cues were used as conditioned stimuli (CS), each of which signaled the delivery of a different unconditioned stimulus (US): sucrose pellet (appetitive outcome), mild foot shock (aversive outcome), or nothing (neutral outcome; **Figure 1A**, left). After 8 sessions of this initial learning phase, the appetitive and aversive associations were reversed, where the CS previously paired with a reward (CS_1_) instead preceded shock delivery and the CS previously paired with a shock (CS_2_) instead preceded reward delivery. Rats were tested on the reversed contingencies for 8 sessions before the associations were re-reversed back to the initial assignments for 8 final sessions. The neutral association (CS-) did not change across the 24 total sessions of FCL (**Figure 1A**, right). Conditioned responding was quantified as the change in the rate of food port head entries during the 5 s CS relative to the 5 s preceding the CS^33^. Rats increased conditioned responding to the food port in response to the initial appetitive CS_1_, but not the aversive CS_2_ or neutral CS-, throughout the initial learning phase (two-way mixed-effects analysis; initial learning phase CS effect: *F*_(1.14, 18.16)_ = 22.71, *p* < 0.0001; session x CS interaction effect: *F*_(2.60, 41.66)_ = 11.81, *p* < 0.0001; **Figure 1B**). When the learned appetitive and aversive associations reversed, rats adapted their behavior by increasing conditioned responding to the newly appetitive CS_2_, and decreased responding during the previously appetitive CS_1_ that became aversive (reversal phase CS effect: *F*_(1.06, 16.98)_ = 12.22, *p* = 0.002; session x CS interaction effect: *F*_(1.75, 26.90)_ = 11.97, *p =* 0.0003; **Figure 1B**). After re-reversal in which CS-US pairings returned to the initial assignments, rats again updated conditioned responding in favor of the appetitive CS_1_ (re-reversal phase CS effect: *F*_(1.13, 15.79)_ = 18.99, *p* = 0.0004; session x CS interaction effect: *F*_(1.73, 23.05)_ = 9.15, *p* = 0.002; **Figure 1B**).

**Figure 1.**
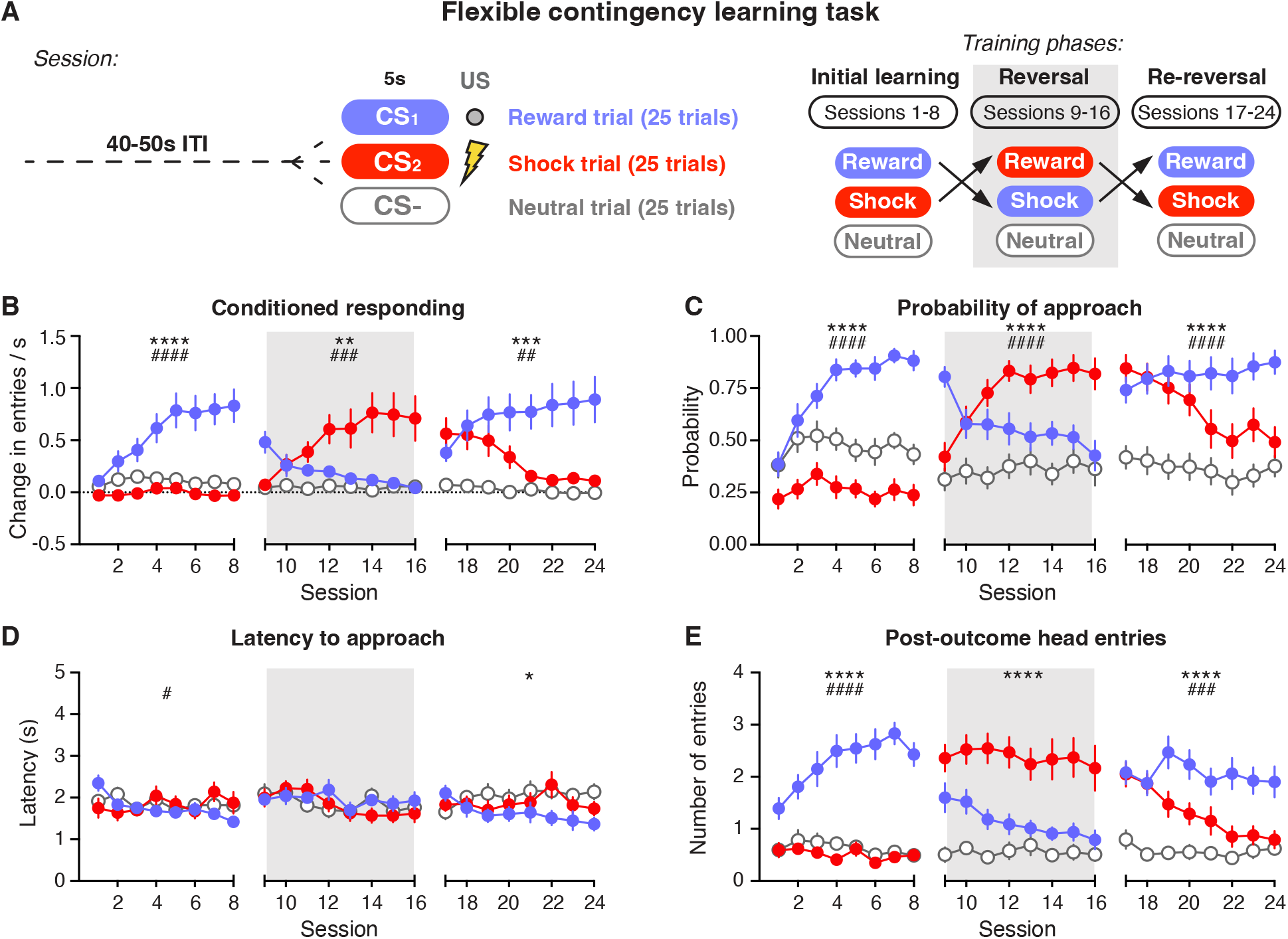
Behavioral responding adapts to reversal of contingencies during FCL. **A**, Schematic for the FCL task. Left, diagram of the three cue-outcome associations presented in every FCL session. Right, chart depicting the initial, reversal, and re-reversal phases of FCL. **B**, Conditioned responding towards CSs initially associated with reward (blue), shock (red) or nothing (gray open circles). **C**, Probability of approaching the food port during CS presentation. **D**, Latency to respond with a head entry into the food port during CS presentation. **E**, Number of food port head entries following US delivery. Gray shading represents reversal period in which appetitive and aversive associations were switched. Data are presented as mean +/-SEM. Asterisks represent main effect of CS; pound signs represent interaction effect with session. #,*p < 0.05; ##,**p < 0.01; ###,*** p < 0.001; ####,**** p < 0.0001.

Conditioned responding adapted when learned associations were reversed. To determine if the rate of responding differed depending on whether the CS was novel or previously paired with an aversive outcome, we compared conditioned responding to the appetitive association between all FCL phases. Rats acquired the appetitive association at similar rates across initial learning, reversal, and re-reversal phases (**Supplemental Figure 1A**). We also examined whether conditioned responding differed between the TH and GAD genotypes, and found no effect of genotype on conditioned responding to the appetitive CS during either initial learning or the reversal phase (**Supplemental Figure 1B**). While the aim of this study was not to examine sex differences, we included both male and female rats and performed further analysis with sex as a factor. There was no effect of sex on conditioned responding to the appetitive CS during either initial learning or the reversal of associations (**Supplemental Figure 1C**). Therefore, data were collapsed across genotype and sex for the remainder of the analyses.

Although subjects reduced rates of conditioned responding to the aversive and neutral CSs throughout FCL (**Figure 1B**), they continued to explore the food port with low levels of approach probability (**Figure 1C**). During the initial learning phase, the appetitive CS_1_ produced the highest probability of approach and the aversive CS_2_ produced the lowest, demonstrating the aversive properties of the shock-paired CS_2_ (initial learning phase CS effect: *F*_(1.61, 25.69)_ = 51.20, *p* < 0.0001; session x CS interaction effect: *F*_(5.33, 85.35)_ = 10.25, *p* < 0.0001; **Figure 1C**). The probability of approach was modified to reflect the new contingencies during reversal (reversal phase CS effect: *F*_(1.99, 31.87)_ = 34.49, *p* < 0.0001; session x CS interaction effect: *F*_(4.94, 75.79)_ = 13.61, *p* < 0.0001), as well as the re-reversal phase (re-reversal phase CS effect: *F*_(1.89, 26.44)_ = 40.29, *p* < 0.0001; session x CS interaction effect: *F*_(4.24, 56.67)_ = 7.14, *p* < 0.0001; **Figure 1C**). When rats approached the food port during a CS, the appetitive CS_1_ also elicited a quicker latency across the initial learning phase (initial learning phase CS effect: *F*_(1.81, 28.98)_ = 0.54, *p* = 0.57; session x CS interaction effect: *F*_(6.85, 104.7)_ = 2.26, *p* = 0.04), and during the second reversal (re-reversal CS effect: *F*_(1.95, 27.32)_ = 4.95, *p =* 0.02; **Figure 1D**). There was no difference in the latency to respond to the appetitive, aversive, or neutral CSs in the reversal phase (**Figure 1D**). Post-outcome head entries increased after the termination of the reward-paired CS_1_ during initial learning (initial learning phase CS effect: *F*_(1.36, 21.79)_ = 81.44, *p* < 0.0001; session x CS interaction effect: *F*_(5.21, 83.38)_ = 7.03, *p* < 0.0001; **Figure 1E**), reversal (reversal phase CS effect: *F*_(1.16, 18.54)_ = 61.81, *p* < 0.0001; session x CS interaction effect: *F*_(4.39, 67.40)_ = 1.60, *p* ***=*** 0.18) and re-reversal (re-reversal phase CS effect: *F*_(1.66, 23.24)_ = 41.51, *p* < 0.0001; session x CS interaction effect: *F*_(4.29, 57.35)_ = 5.40, *p* = 0.0007; **Figure 1E**). These findings illustrate that rats learn to distinguish between simultaneously acquired appetitive, aversive, and neutral cues, and adapt their behavior when contingencies update.

### Dissociable VTA dopamine and GABA responses dynamically adapt during learning and reversal of contingencies

VTA dopamine and GABA neurons are implicated in associative learning, but their distinct roles in flexible updating of cue-outcome associations is not well understood. We used fiber photometry to measure cue- and outcome-elicited calcium responses in VTA dopamine and GABA neuron populations throughout FCL. To record from the VTA dopamine neuron population, TH::cre rats expressed GCaMP6f in TH+ cells in the VTA, and fiber placement was centered above the virus injection (**Figure 2A**). Differences in neural calcium activity between the first and last session of each FCL phase were assessed using a permutation-based approach^35,38^. In the first session (session 1), VTA dopamine calcium activity increased in response to reward but not the appetitive CS_1_ predicting reward. By the end of the initial learning phase (session 8), a large phasic increase in response to the appetitive predictive CS_1_ developed (**Figure 2B**). The dopamine response to the aversive CS_2_ or neutral CS-did not significantly change across initial sessions (**Figure 2B**). By session 8, however, dopamine population calcium activity was lower during the aversive CS_2_ compared to the neutral CS-(**Supplemental Figure 2A**). When the appetitive and aversive associations reversed (session 9), the dopamine population first displayed a prediction error-like response: decreased activity to the unexpected shock and increased activity to the unexpected reward (**Figure 2C**; **Supplemental Figure 2B; Supplemental Figure 3A**). By the end of the reversal phase (session 16), the dopamine population had adjusted to the new contingencies and displayed a similar response profile to the contingencies as before the reversal (**Figure 2C**; **Supplemental Figure 4A-B**). During re-reversal, the dopamine population followed a similar pattern by initially responding in a prediction error-like manner (session 17) and adapted by the end of the phase (session 24; **Figure 2D**; **Supplemental Figure 2C; Supplemental Figure 3B**; **Supplemental Figure 4A-B**).

**Figure 2.**
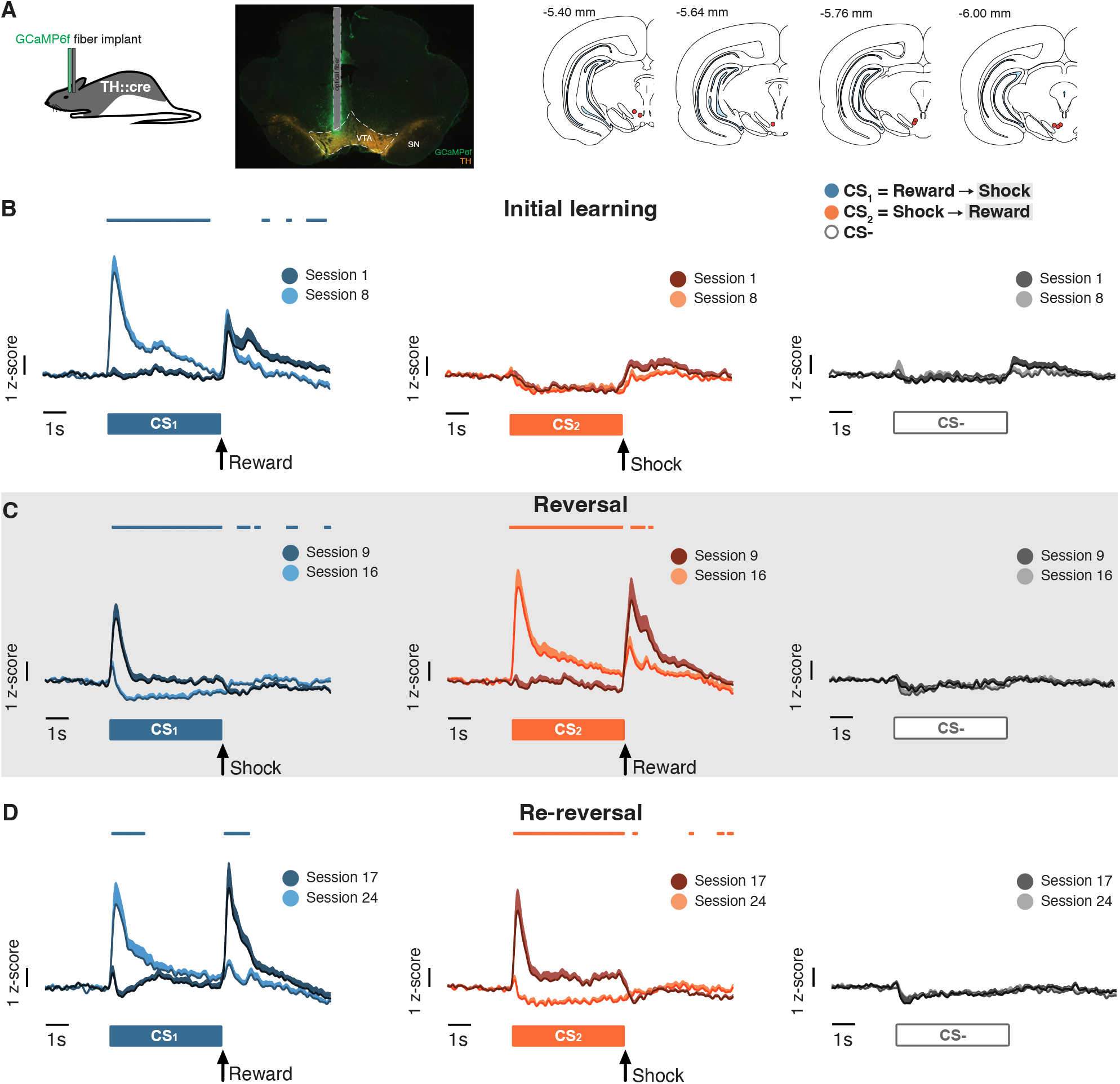
VTA dopamine signaling encodes reward association across FCL phases. **A**, Left, GCaMP6f was expressed in dopamine neurons of TH::cre rats. Right, optic fibers were placed above viral expression in the VTA. **B**, Average VTA dopamine population calcium activity towards CSs associated with reward (teal, left), shock (orange, middle) or nothing (gray, right) during the first (Session 1) and last (Session 8) sessions of the initial learning phase. The first session of each phase is depicted in darker shades, the last session of each phase is depicted in lighter shades. Colored lines above each trace represent a significant difference between the first and last session of the phase detected via permutation test. **C**, Average calcium activity towards CSs associated with shock (teal, left), reward (orange, middle) or nothing (gray, right) during the first (Session 9) and last (Session 16) sessions of the reversal phase. **D**, Average calcium activity towards CSs associated with reward (teal, left), shock (orange, middle) or nothing (gray, right) during the first (Session 17) and last (Session 24) sessions of the re-reversal phase. Data are presented as mean + SEM. Vertical scale bars indicate 1 z-score. Horizontal scale bars indicate 1 second.

To record from the VTA GABA neuron population, we used fiber photometry in GAD::cre rats expressing GCaMP6f in GAD+ cells (**Figure 3A**). In session 1, the GABA population displayed a phasic increase in response to all three CSs as well as both shock and reward USs (**Figure 3B**; **Supplemental Figure 2D**). Despite subjects successfully learning the contingencies, by the end of the initial phase GABA activity in response to these events, including the neutral CS, remained the same (**Figure 1**; **Figure 3B**; **Supplemental Figure 1**; **Supplemental Figure 2D)**. The only exception was that the GABA response to appetitive CS_1_ increased after learning (**Figure 3B**; **Supplemental Figure 2D**). When the appetitive and aversive associations reversed, the GABA population also displayed a prediction error-like response in session 9 by decreasing activity to the unexpected shock and increasing activity to the unexpected reward (**Figure 3C**; **Supplemental Figure 2E; Supplemental Figure 3C**). By the end of reversal learning, GABA responses were nearly identical to the last session of initial learning (**Figure 3C**; **Supplemental Figure 4C-D**). During the second reversal, the GABA population again showed a prediction error-like response and adapted by the end of the re-reversal phase (**Figure 3D**; **Supplemental Figure 2F; Supplemental Figure 3D**; **Supplemental Figure 4C-D**). The GABA response to the neutral CS remained unchanged even after extensive training.

**Figure 3.**
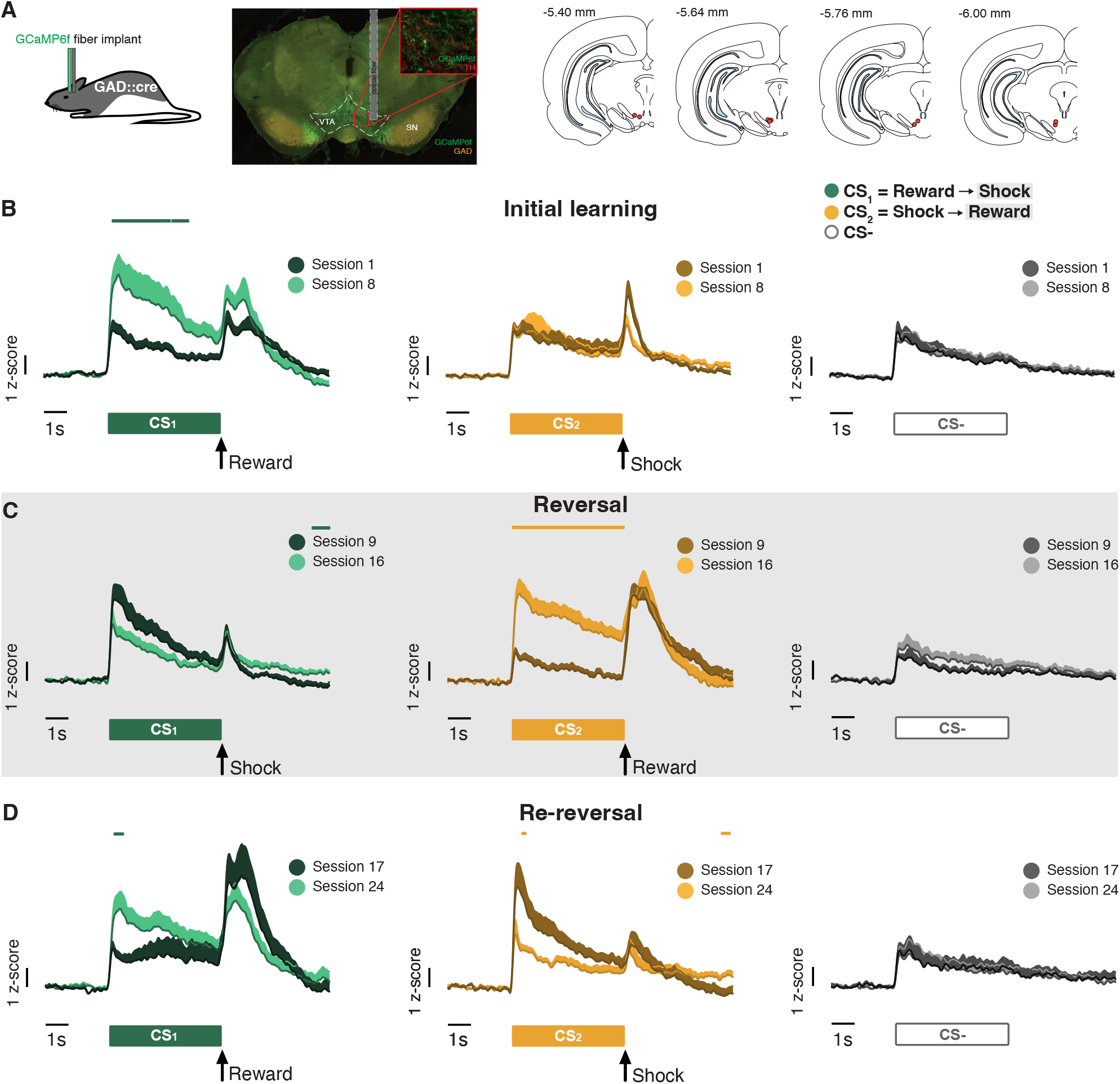
VTA GABA population responds to all associations across FCL phases. **A**, Left, GCaMP6f was expressed in GABA neurons of GAD::cre rats, and there was no co-expression with TH neurons (inset). Right, optic fibers were placed above viral expression in the VTA. **B**, Average VTA GABA population calcium activity towards CSs associated with reward (green, left), shock (yellow, middle) or nothing (gray, right) during the first (Session 1) and last (Session 8) sessions of the initial learning phase. The first session of each phase is depicted in darker shades, the last session of each phase is depicted in lighter shades. Colored lines above each trace represent a significant difference between the first and last session of the phase detected via permutation test. **C**, Average calcium activity towards CSs associated with shock (green, left), reward (yellow, middle) or nothing (purple, gray) during the first (Session 9) and last (Session 16) sessions of the reversal phase. **D**, Average calcium activity towards CSs associated with reward (green, left), shock (yellow, middle) or nothing (gray, right) during the first (Session 17) and last (Session 24) sessions of the re-reversal phase. Data are presented as mean + SEM. Vertical scale bars indicate 1 z-score. Horizontal scale bars indicate 1 second.

When a CS was contingent on an outcome, neural calcium activity of both dopamine and GABA populations dynamically changed between the first and last session of each phase in FCL (**Figures 2-3**). To quantify the flexible progression of CS-evoked dopamine and GABA responses in different FCL phases, we averaged activity during the full 5 s CS presentation, as well the early (first 2 s) and late (last 3 s) periods of CS presentation (**Figure 4A-D**) over all sessions. When considering the full CS response in the initial learning phase, VTA dopamine activity was largest to the appetitive CS_1_ compared to the aversive and neutral CSs (initial learning phase CS effect: *F*_(1.82, 12.79)_ = 43.07, *p* < 0.0001; session x CS interaction effect: *F*_(3.28, 21.52)_ = 5.42, *p* = 0.005; **Figure 4A**). By the end of the initial learning phase, dopamine responses to the shock-predictive CS_2_ were lower than to the neutral CS-. This pattern of responding flexibly updated to the new contingencies during the reversal phase (reversal phase CS effect: *F*_(1.19, 8.32)_ = 21.34, *p* = 0.001; session x CS interaction effect: *F*_(2.83, 17.96)_ = 10.30, *p* = 0.0004) as well as during the re-reversal phase (re-reversal phase CS effect: *F*_(1.33, 9.33)_ = 22.81, *p* = 0.0005; session x CS interaction effect: *F*_(2.15, 11.81)_ = 10.61, *p* = 0.002; **Figure 4A**). In contrast to VTA dopamine, VTA GABA calcium activity ubiquitously increased to all CSs across the initial learning phase (initial learning phase CS effect: *F*_(1.30, 10.37)_ = 3.77, *p* = 0.07; session x CS interaction effect: *F*_(2.68, 20.29)_ = 2.15, *p* = 0.13; **Figure 4B**). Reversing the learned contingencies, however, increased GABA activity to the new appetitive CS_2_ compared to the other CSs (reversal phase CS effect: *F*_(1.44, 11.50)_ = 6.00, *p* = 0.02; session x CS interaction effect: *F*_(2.34, 16.71)_ = 7.82, *p* = 0.003; **Figure 4B**). This response pattern from the GABA population persisted into the re-reversal phase with an interaction effect between session and CS (re-reversal phase CS effect: *F*_(1.03, 7.23)_ = 4.32, *p* = 0.07; session x CS interaction effect: *F*_(2.60, 13.20)_ = 4.31, *p* = 0.03; **Figure 4B**).

**Figure 4.**
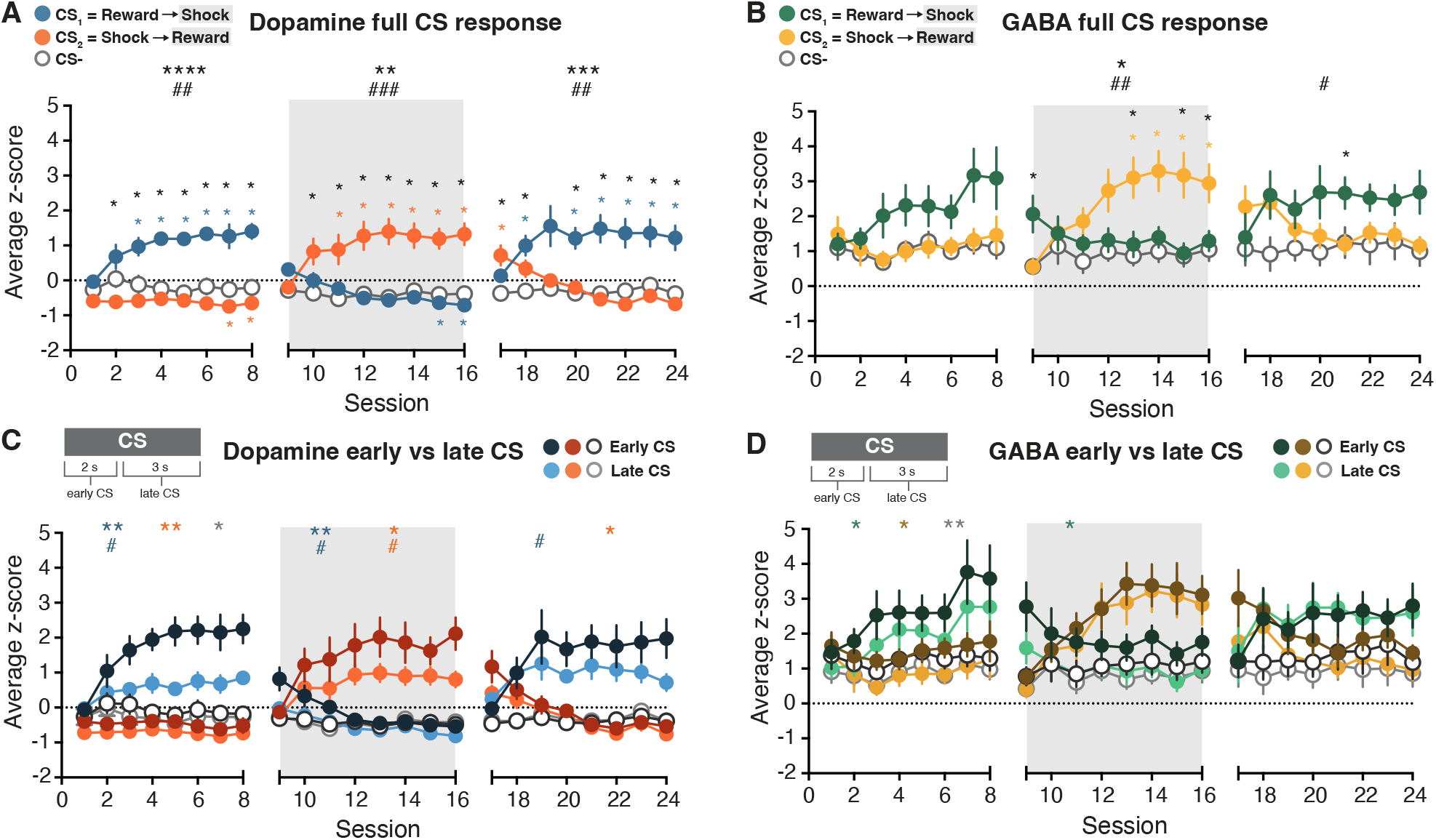
Quantification of VTA dopamine and GABA responses evoked by full, early or late periods of CS presentation across FCL sessions. **A**,**B**, Dopamine (A) and GABA (B) average calcium activity during the full 5 s CS presentation. Large asterisks above the graph represent main effect of CS, pound signs represent interaction effect with session. Small asterisks within graph represent post-hoc comparisons between CSs: black asterisks compare CS_1_ and CS_2_; teal (A) and green (B) asterisks compare CS_1_ and CS-; orange (A) and yellow (B) asterisks compare CS_2_ and CS-. **C**,**D**, Dopamine (C) and GABA (D) average calcium activity during the early period (first 2 s; darker shades) or the late period (last 3 s; lighter shades) of CS presentation. Data are presented as mean +/-SEM. Asterisks represent main effect between early and late CS, pound signs represent interaction effect with session. #,*p < 0.05; ##,**p < 0.01; ###,*** p < 0.001; ####,**** p < 0.0001.

Regardless of task phase or predicted outcome valence, throughout FCL CS-evoked GABA activity was higher than dopamine activity (**Supplemental Figure 5A**). Whereas VTA dopamine calcium activity displayed a phasic increase at CS onset that diminished towards baseline by the end of the CS (in the case of the appetitive association), GABA calcium activity remained amplified throughout the CS (see **Figures 2-3**). We therefore quantified differences between the early CS period (0-2 s after CS onset) and the late CS period (2-5 s after CS onset; **Figure 4C-D**). In the initial learning phase, every CS produced a larger response during early CS compared to late CS in both the dopamine population (initial learning phase CS_1_ early v late CS effect: *F*_(1, 7)_ = 21.87, *p* = 0.002; CS_1_ session x early v late CS interaction effect: *F*_(2.45, 15.78)_ = 5.74, *p* = 0.01; CS_2_ early v late CS effect: *F*_(1, 7)_ = 19.91, *p* = 0.003; CS-early v late CS effect: *F*_(1, 7)_ = 10.04, *p* = 0.02; **Figure 4C**) and the GABA population (CS_1_ early v late CS effect: *F*_(1, 8)_ = 7.63, *p* = 0.02; CS_2_ early v late CS effect: *F*_(1, 8)_ = 9.71, *p* = 0.01; CS-early v late CS effect: *F*_(1, 8)_ = 14.35, *p* = 0.005; **Figure 4D**). The early response to the appetitive and aversive CSs continued in the dopamine population throughout reversal and re-reversal phases (reversal phase CS_1_ early v late CS effect: *F*_(1, 7)_ = 13.36, *p* = 0.008; CS_1_ session x early v late CS interaction effect: *F*_(2.49, 15.29)_ = 5.00, *p* = 0.02; CS_2_ early v late CS effect: *F*_(1, 7)_ = 8.56, *p* = 0.02; CS_2_ session x early v late CS interaction effect: *F*_(2.53, 15.53)_ = 3.63, *p* = 0.04; re-reversal phase CS_1_ session x early v late CS interaction effect: *F*_(1.69, 8.46)_ = 7.82, *p* = 0.01; CS_2_ early v late CS effect: *F*_(1, 7)_ = 9.07, *p* = 0.02; **Figure 4C**). The reversal phase also increased reward-evoked US responses in dopamine and GABA populations (**Supplemental Figure 5B-E**). However, reversing the contingencies eliminated differences in GABA activity between early and late CS responses for the new appetitive CS_2_ and neutral CS-, and for all CSs during re-reversal (**Figure 4D**). Thus, the CS-evoked activity during the reversal phase was encoded differently by GABA, and not dopamine, populations. The GABA response to the appetitive CS significantly increased during reversal, and differences between early and late CS-evoked activity were no longer significant.

### Reversing learned associations increases within-session activity in the VTA GABA population

Dopamine and GABA calcium activity changed over FCL sessions, reflecting learning by both populations across days (**Figures 2-4**). To determine the temporal dynamics of this change, we parsed sessions into 5-trial bins and compared full CS responses between the first bin and the last bin in each session (**Figure 5A-B**; **Supplemental Figure 6; and equivalent analysis of behavioral responding in Supplemental Figure 7**). In all FCL phases, both dopamine and GABA populations exhibited within-session increases in CS-evoked calcium activity. However, there was a stark difference in the temporal progression of within-session activity between dopamine and GABA populations. In particular, CS-evoked dopamine activity increased to the appetitive association within the first several sessions of each FCL phase, whereas CS-evoked GABA within-session activity increased to both appetitive and aversive associations throughout the FCL phases (**Supplemental Figure 6A**,**C**). To quantify these differences, we averaged the binned activity between early (first 4) and late (last 4) sessions of each FCL phase (**Figure 5C**).

**Figure 5.**
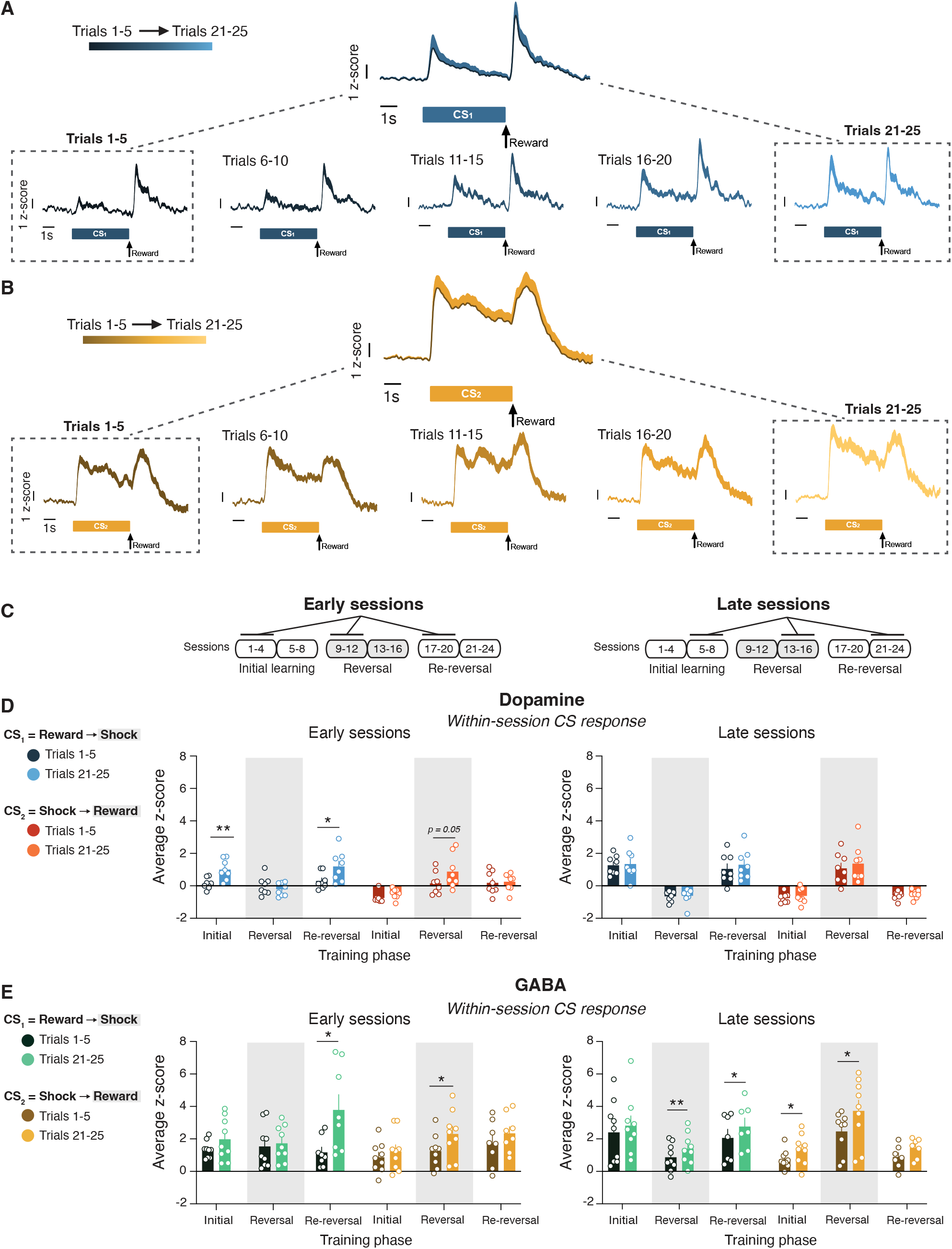
Within-session changes in CS-evoked calcium activity in VTA dopamine and GABA populations differ between early and late sessions in each FCL phase. **A**, Representative trace of within-session changes in calcium activity during an early phase session. Example trace of VTA dopamine calcium activity during a single session (Session 2) to the appetitive CS_1_ portioned into 5-trial bins. Dotted rectangles represent first and last 5-trial bins from the session. **B**, Representative trace of within-session changes in calcium activity during a late phase session. Example trace of VTA GABA calcium activity during a single session (Session 14) to the appetitive CS_2_ portioned into 5-trial bins. **C**, Schematic depicting early (first 4 sessions; left) and late (last 4 sessions; right) sessions of each FCL phase. **D**, Dopamine within-session calcium activity during the 5 s CS period for the appetitive and aversive associations in early (left) and late (right) sessions. Sessions were split into 5-trial bins, and bins were averaged across sessions. The color intensity represents the beginning of the session (Trials 1-5, dark shades) or the end of the session (Trials 21-25, light shades). **E**, GABA within-session calcium activity during the 5 s CS period for the appetitive and aversive associations in early (left) and late (right) sessions. Data are presented as mean +/-SEM, with individual subject points included. Asterisks represent post-hoc comparisons between trial bins for each CS. *p < 0.05; **p < 0.01.

The temporal pattern of within-session responses to the appetitive CS were different between dopamine and GABA populations. Dopamine calcium activity increased to the reward-predictive CS within only the early sessions of every FCL phase (dopamine early sessions CS_1_ trial effect: *F*_(1, 7)_ = 28.22, *p* = 0.001; FCL phase x CS_1_ trial interaction effect: *F*_(1.96, 13.69)_ = 9.95, *p* = 0.002; CS_2_ trial effect: *F*_(1, 7)_ = 7.75, *p* = 0.03; FCL phase x CS_2_ trial interaction effect: *F*_(1.35, 9.43)_ = 4.78, *p* < 0.05; **Figure 5D**). In contrast, the GABA population did not exhibit within-session changes in activity to the appetitive association during the initial learning phase (**Figure 5E**).

Reversing the learned contingencies, however, elicited increased within-session GABA activity to the appetitive association during both early and late sessions, an effect that persisted into the re-reversal phase (GABA early sessions CS_1_ trial effect: *F*_(1, 8)_ = 20.33, *p* = 0.002; FCL phase x CS_1_ trial interaction effect: *F*_(1.33, 9.28)_ = 8.22, *p* = 0.01; CS_2_ trial effect: *F*_(1, 8)_ = 15.86, *p* = 0.004; late sessions CS_1_ trial effect: *F*_(1, 8)_ = 14.37, *p* = 0.005; CS_2_ trial effect: *F*_(1, 8)_ = 29.74, *p* = 0.0006; **Figure 5E**). Furthermore, only the GABA population exhibited increased within-session activity to the aversive association, which occurred during late sessions of the initial learning and reversal phases (**Figure 5E**). Changes in dopamine and GABA responses to the neutral CS-, as well as to the reward and shock USs, remained minimal within FCL sessions (**Supplemental Figure 6**). Within-session analysis of behavior indicates that conditioned responding increased to the appetitive association during both early and late sessions throughout FCL phases, implicating both dopamine and GABA within-session increased activity in modulating behavioral responding (**Supplemental Figure 7**). Thus while within-session signaling is modulated in both VTA dopamine and GABA populations, it is dissociable through distinct temporal and valence-specific response profiles.

## Discussion

A large number of studies have characterized the activity of VTA dopamine neurons during reward learning^5-15^, whereas the role of GABA activity has been primarily limited to its modulation of dopamine function^19-24^. Some prior research, however, has shown that VTA GABA neurons elicit responses that are distinct from dopamine neurons^8,29,30,32^, suggesting that VTA GABA and dopamine populations may have independent roles in encoding associative learning. Here, we compared VTA dopamine and GABA calcium activity using fiber photometry during a flexible contingency learning paradigm that assessed initial learning and reversal of appetitive and aversive associations acquired simultaneously. The initial acquisition of cue-outcome associations elicited responses from both dopamine and GABA VTA populations. Reversing the learned reward and punishment contingencies, however, selectively influenced GABA population responses to the reward-predictive cue, by prolonging increased calcium activity both within and across sessions. These findings reveal that the VTA GABA population is critically modulated when learned contingencies update, and further suggests that dopamine and GABA neurons are independently recruited during learning of appetitive and aversive associations.

The ability to flexibly update responding when learned contingencies change has been examined across species through reversal learning, latent inhibition, and counterconditioning paradigms^40-42^. Reversal learning typically involves switching between two appetitive outcomes, and latent inhibition comprises overriding a neutral stimulus with an association that has valence. Counterconditioning paradigms, which employ reversals between appetitive and aversive associations, examine each valence reversal independently^43,44^. Our flexible contingency learning task combines features of these paradigms to demonstrate that rats adapt to the valence reversal of reward and punishment associations experienced concurrently^31,32^. Our current findings extend these observations to both male and female rats, and we demonstrate rats also re-reverse their behavior when associations are returned to their initial contingencies. We did not identify sex differences in behavioral responding, potentially due to the strain used in this study^33,45^. Pre-exposure to cues associated with a neutral outcome (such as in latent inhibition) or an aversive outcome (such as in counterconditioning) has been found to dampen behavioral responding when the contingency is subsequently shifted to an appetitive outcome^41-43,46,47^. However, in our task where multiple cue-outcome associations are experienced in the same session, we find that rats learn to associate the previously aversive predictive cue with a rewarding outcome at the same rate as the initial acquisition of the appetitive association. This suggests that a more complex environment may facilitate behavioral flexibility.

We observed distinct calcium influx responses between dopamine and GABA neurons in the VTA to appetitive, aversive, and neutral associations. On a population level, VTA dopamine neuron calcium activity increased to the appetitive association and decreased to the aversive association, with minimal response to the neutral cue. This is consistent with the response profile for the majority of individual VTA dopamine neurons examined previously^5,8,10,11,16,17,31^. In contrast, VTA GABA population calcium activity increased in response to all salient stimuli. Initially this increased response was uniform, but after additional exposure the appetitive cue evoked a more pronounced increase in GABA activity relative to the aversive and neutral cues. Although changes in calcium activity measured with fiber photometry do not necessarily reflect spiking activity^48^ (but see^49^), our findings are consistent with previously measured changes in spike rate of VTA dopamine and GABA neurons^8,31^. Furthermore, this approach allowed us to observe responses to the shock outcome, which causes noise in electrophysiological recordings. Both shock and reward outcomes as well as all predictive cues, including the neutral cue regardless of the extensive exposure to this association, elicited increases in VTA GABA calcium activity, implicating this population in salience signaling^50^. When the contingencies were reversed, both VTA dopamine and GABA calcium activity exhibited a reward prediction error-like response. The purpose of prediction error responses in dopamine neurons have been well-studied as a teaching signal for adaptive learning^5,6,8-10,51,52^. Limited prior research has also identified increased VTA GABA firing to the delivery of an unexpected reward^53^, which may function to convey outcome value to downstream targets such as the ventral pallidum^54^.

While both dopamine and GABA population calcium activity increased to the appetitive association, activation of these populations produces different effects on reward-based behavioral responding. Optogenetic studies have shown that activating VTA dopamine neurons during either an appetitive cue or reward delivery enhances cued reward seeking^51,55^. In contrast, activation of the VTA GABA population prior to or during reward delivery respectively decreases anticipatory conditioned responding and reward consumption^22,53^. Nevertheless, fiber photometry and optogenetic approaches used in vivo are unable to disentangle potential heterogeneity between individual neurons. We appreciate that both dopamine and GABA cell groups in the VTA can display diverse responses to stimuli that can depend on their anatomical location or projection targets^3,18,30,56^. Our observation that within-session changes in cue-evoked GABA activity occur throughout training suggests that we likely recorded signals from both local and long-range projecting GABA neurons, which may individually be critical at separate stages of training. Elevated local VTA GABA activity could serve as a salience or prediction signal to dopamine neurons during initial learning^50,53^, while increased activity in long-range GABA projections to cholinergic interneurons in the ventral striatum could facilitate cue discrimination during reversal sessions^25^. Understanding the multifaceted role of VTA GABAergic activity during associative learning will be essential for future research.

In conclusion, we identify distinct patterns of responses between VTA dopamine and GABA populations during appetitive and aversive associative learning and contingency reversal. A critical observation was that VTA GABA neuron population calcium activity is selectively amplified by contingency reversal. This finding supports a role for GABA neurons in behavioral flexibility. Previous research examining flexible behavior focus on how cortical, striatal, and amygdalar regions mediate the reversal of learned contingencies^40-42^, which all send projections to the VTA^29,57-60^. Therefore, it will be critical to determine the impact of the afferent projections on VTA dopamine and GABA signaling when learned contingencies are updated.

## Author contributions

M.J.L. and B.M. designed research; M.J.L. performed research, analyzed data, and wrote the first draft of the paper; M.J.L. and B.M. edited the paper; M.J.L. and B.M. wrote the paper.

## Competing interests

The authors report no conflicts of interest.

## Acknowledgements

This work was supported by Public Health Service awards from the National Institute of Mental Health (Grant No. R01-MH048404 [to BM]) and the National Institute on Drug Abuse (Grant No. F32-DA 060070 [to MJL] and T32-DA007262 [to MJL]). We thank Dr. David Jacobs, Dr. Michelle Kielhold, and Dr. Alejandro Torrado Pacheco for thoughtful commentary on the manuscript. We thank Alina Bogachuk for technical assistance.

## Data Availability

The datasets presented in the current study are available from the corresponding author upon reasonable request.

## Notes

### Competing Interest Statement

The authors have declared no competing interest.

