## Supplementary files for "Reward and punishment contingency shifting reveals distinct roles for VTA dopamine and GABA neurons in behavioral flexibility"

**Lefner and Moghaddam supplementary files**

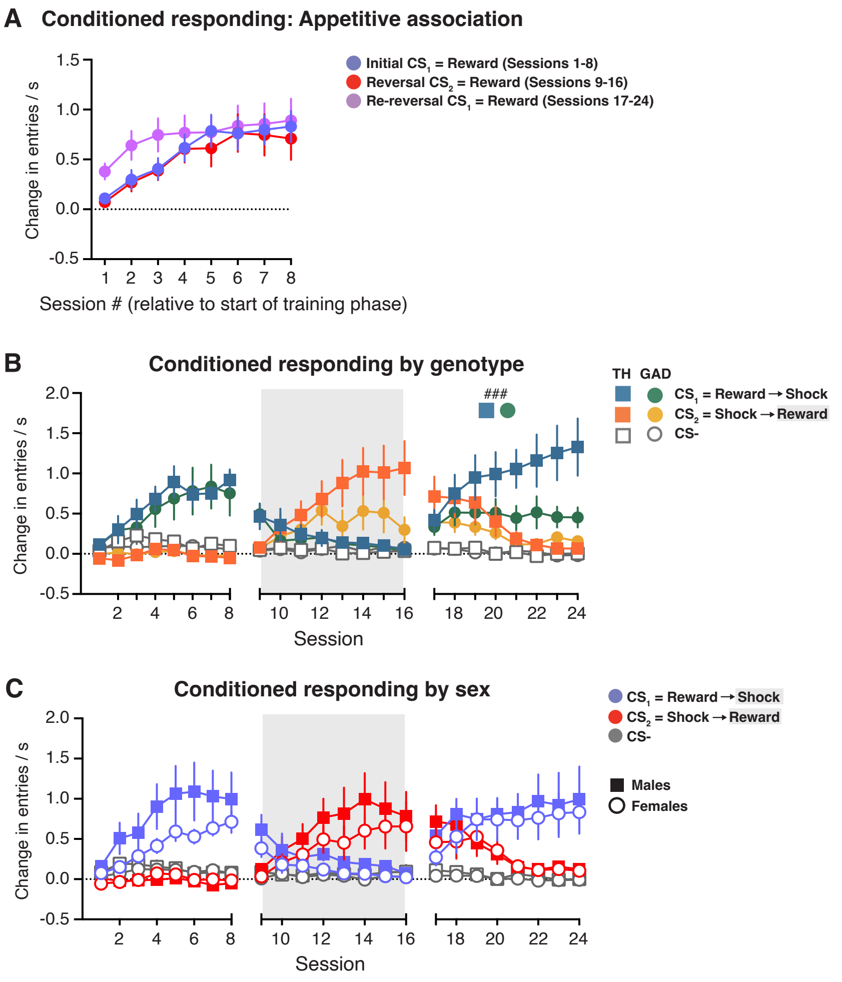


*Supplemental Figure 1*. Additional analysis of behavioral and neural responses during FCL. **A**, Conditioned responding to the appetitive association compared between initial learning, reversal, and re-reversal phases. **B**, Conditioned responding throughout FCL compared between TH (squares) and GAD (circles) rats. **C**, Conditioned responding throughout FCL compared between male (closed squares) and female (open circles) rats. Data are presented as mean +/- SEM. ### $p < 0.001$.

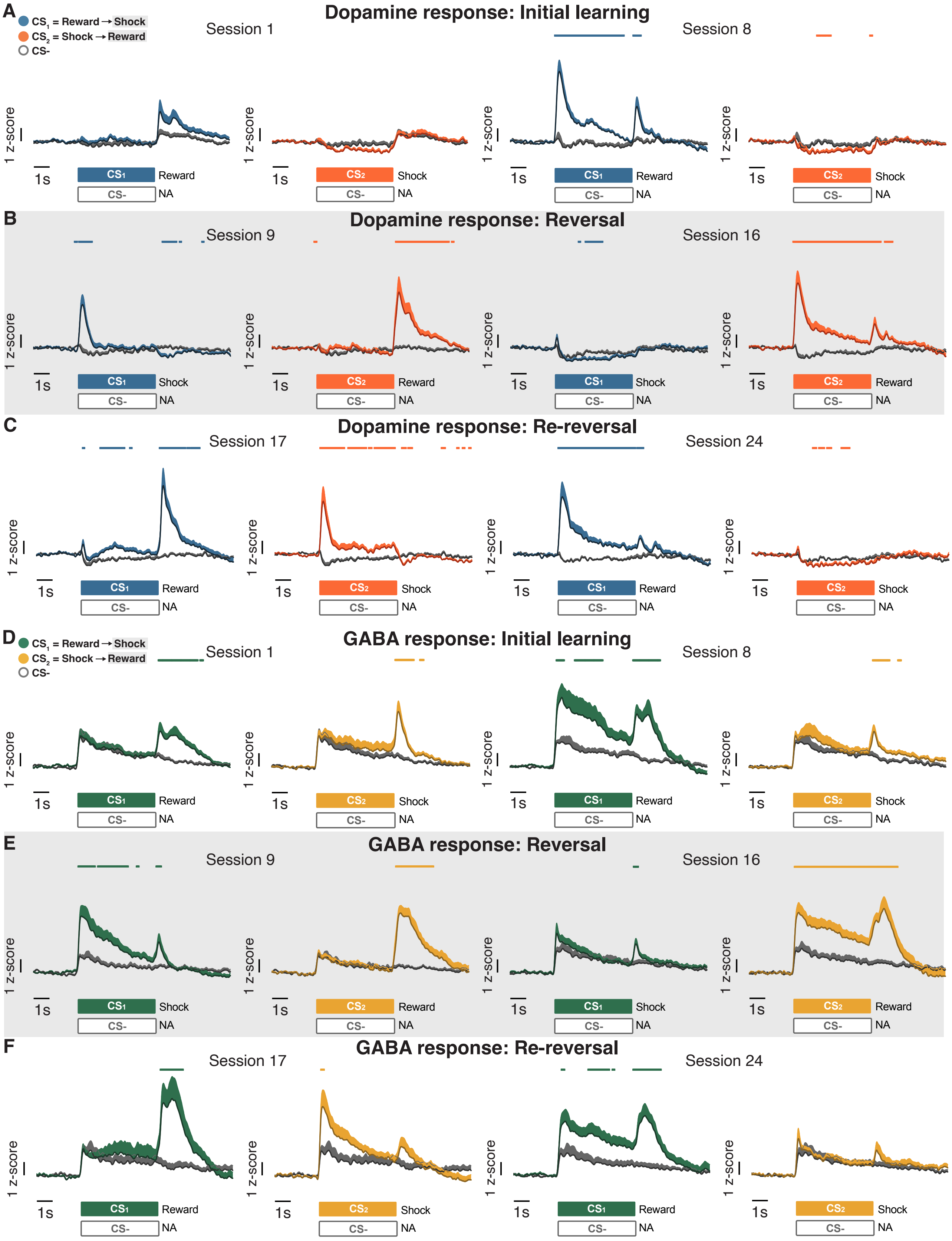


*Supplemental Figure 2*. VTA dopamine and GABA calcium activity to appetitive and aversive CSs in comparison to the neutral CS-. **A**, Average VTA dopamine calcium activity during first (Session 1) and last (Session 8) sessions of initial learning. Colored lines above each trace represent a significant difference between the reward- or shock-paired CSs compared to the neutral CS detected via permutation test. **B**, Average VTA dopamine calcium activity during first (Session 9) and last (Session 16) sessions of reversal.

**C**, Average VTA dopamine calcium activity during first (Session 17) and last (Session 24) sessions of re-reversal. **D**, Average VTA GABA calcium activity during first (Session 1) and last (Session 8) sessions of initial learning. **E**, Average VTA GABA calcium activity during first (Session 8) and last (Session 16) sessions of reversal. **F**, Average VTA GABA calcium activity during first (Session 17) and last (Session 24) sessions of re-reversal. Data are presented as mean + SEM.

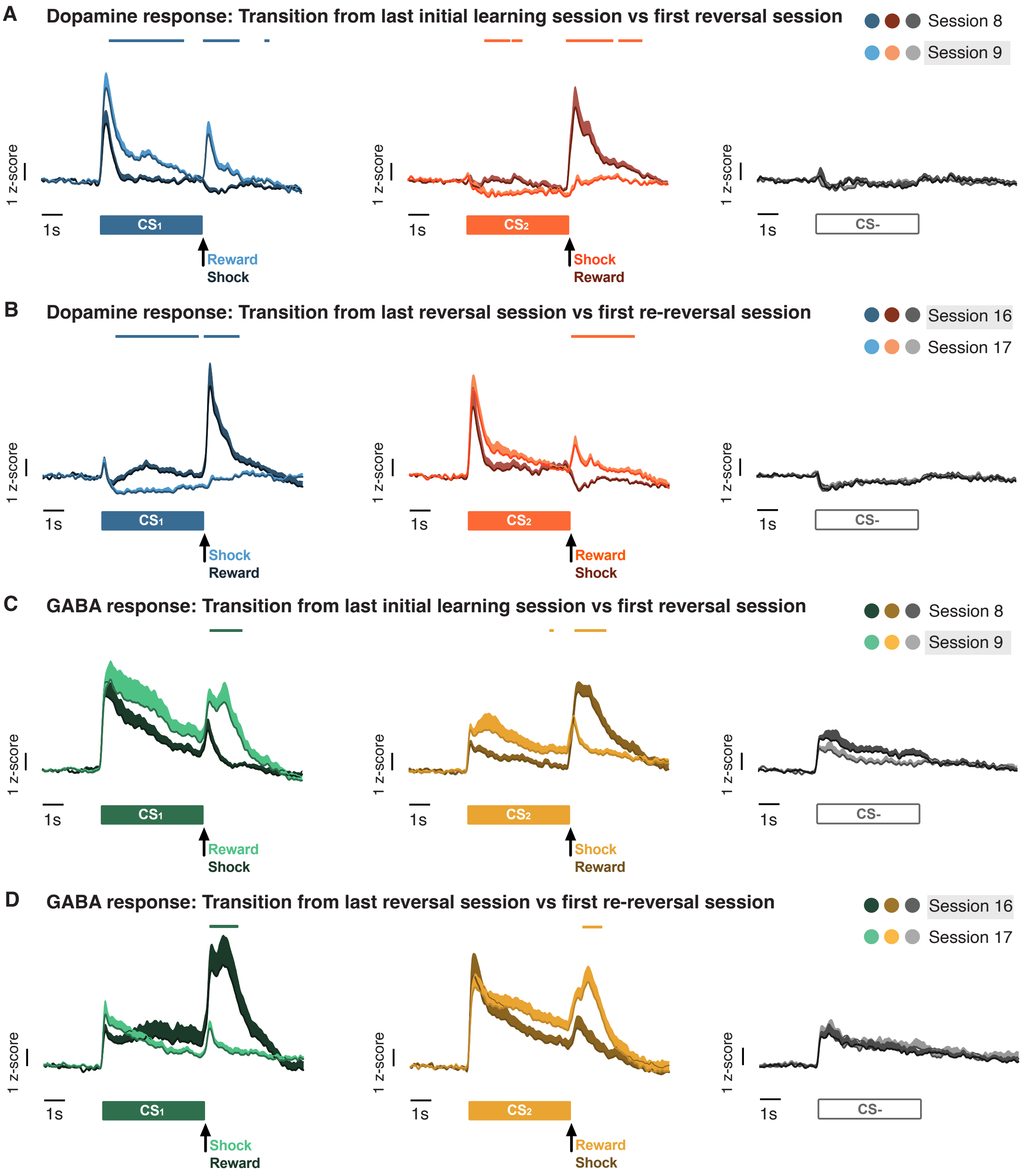


*Supplemental Figure 3*. VTA calcium activity compared between last session of prior phase and first reversal session. **A**, Average VTA dopamine calcium activity between the last initial learning session (Session 8) and the first reversal session (Session 9). The last session of a phase is depicted in lighter shades, compared to first session of the subsequent phase depicted in darker shades. Colored lines above each trace represent a significant difference between the last session of the prior phase and the first session of the subsequent phase detected via permutation test. **B**, Average VTA dopamine calcium activity between the last reversal session (Session 16) and the first re-reversal session (Session 17). **C**, Average VTA GABA calcium activity between the last initial learning session (Session 8) and the first reversal session (Session 9). **D**, Average VTA GABA calcium activity between the last reversal session (Session 16) and the first re-reversal session (Session 17). Data are presented as mean + SEM.

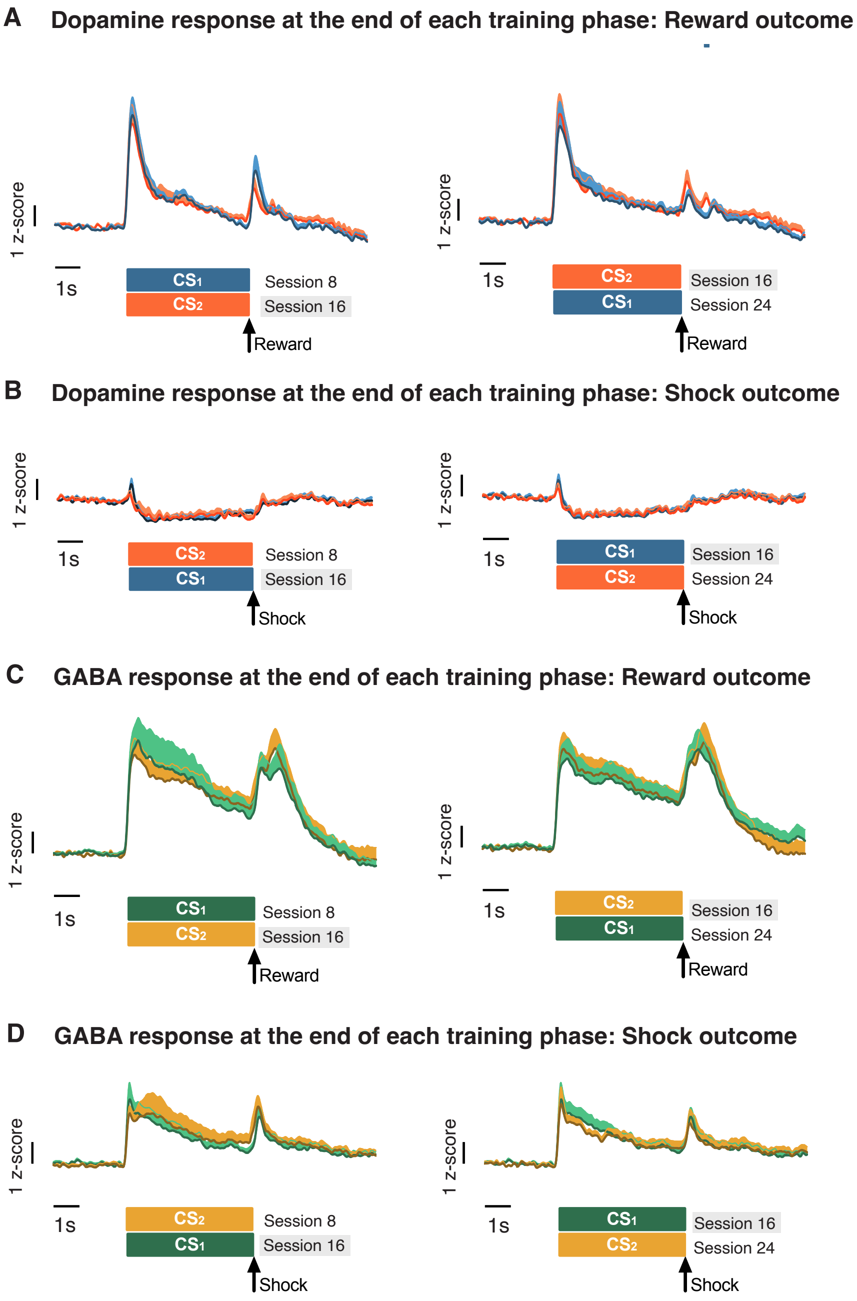


*Supplemental Figure 4*. VTA calcium activity compared between last session of the prior phase and last session of current phase. **A**, Average VTA dopamine calcium activity to CSs predicting the reward US. Left, reward US comparison between the last initial session (Session 8) and the last reversal session (Session 16). Right, reward US comparison between the last reversal session (Session 16) and the last re-reversal session (Session 24). Colored lines above each trace represent a significant difference between the last session of the prior phase and the last session of the subsequent phase detected via permutation test. **B**, Average VTA dopamine calcium activity to CSs predicting the shock US. Left, shock US comparison between the last initial session (Session 8) and the last reversal session (Session 16). Right, shock US comparison between the last reversal session (Session 16) and the last re-reversal session (Session 24). **C**, Average VTA GABA calcium activity to CSs predicting the reward US. Left, reward US comparison between the last initial session (Session 8) and the last reversal session (Session 16). Right, reward US comparison between the last reversal session (Session 16) and the last re-reversal session (Session 24). **D**, Average VTA GABA calcium activity to CSs predicting the shock US. Left, shock US comparison between the last initial session (Session 8) and the last reversal session (Session 16).

Right, shock US comparison between the last reversal session (Session 16) and the last re-reversal session (Session 24). Data are presented as mean + SEM.

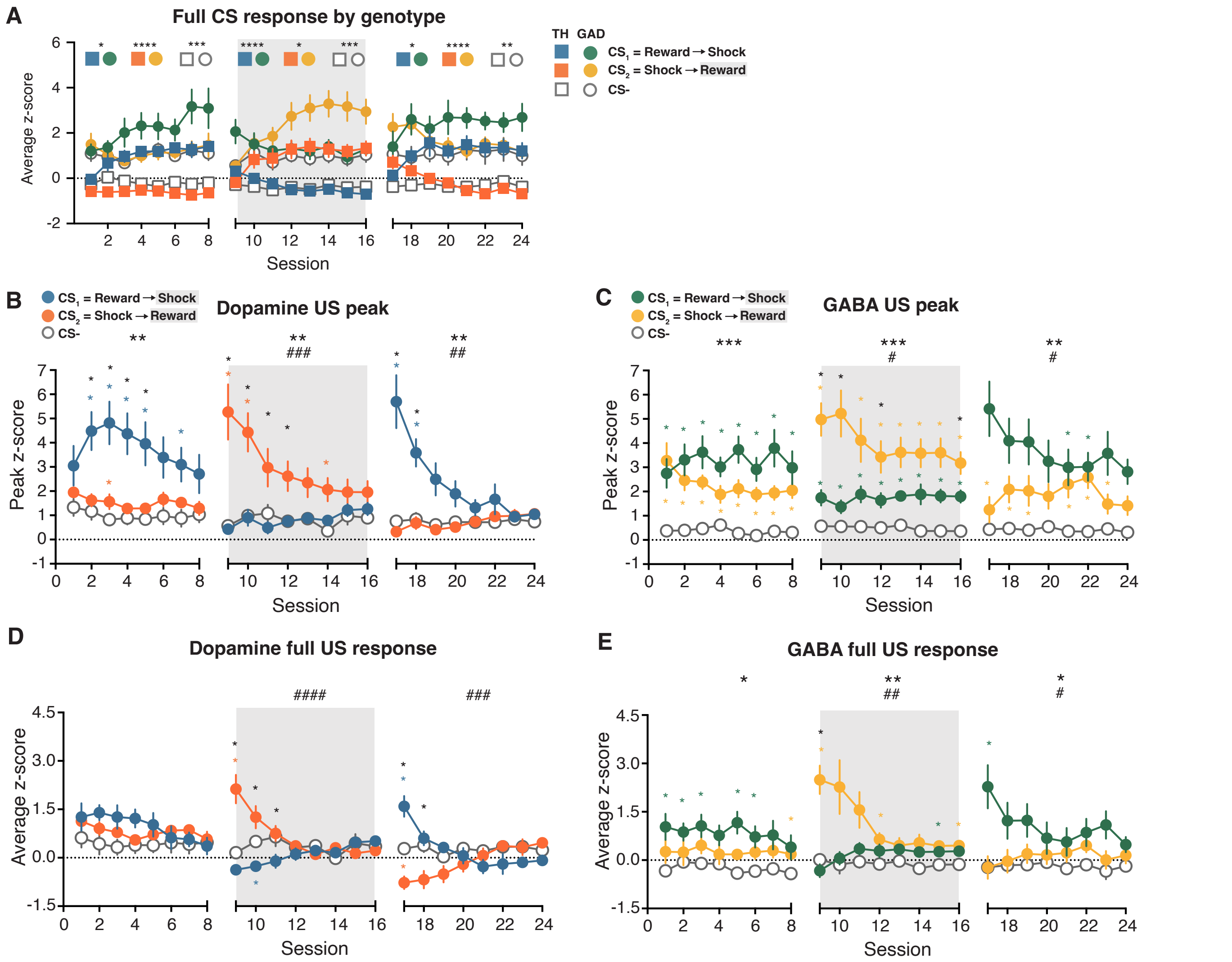


*Supplemental Figure 5*. Quantified VTA calcium activity. **A**, Average calcium activity during the 5s CS compared between TH (squares) and GAD (circles) rats throughout FCL. **B**,**C**, Dopamine (B) and GABA (C) average calcium activity during the 3 s following US delivery, relative to the 0.5 s prior. **D**,**E**, Dopamine (D) and GABA (E) peak value during the 3 s following US delivery, relative to the 0.5 s prior. Data are presented as mean +/- SEM. Large asterisks above graphs B-E represent main effect of US, pound signs represent interaction effect with session. Small asterisks within graph represent post-hoc comparisons between USs: black asterisks compare $US_1$ and $US_2$; teal (Dopamine) and green (GABA) asterisks compare $US_1$ and US-; orange (TH) and yellow (GAD) asterisks compare $US_2$ and US-. #,*p < 0.05; ##,**p < 0.01; ###,*** p < 0.001; ####,**** p < 0.0001.

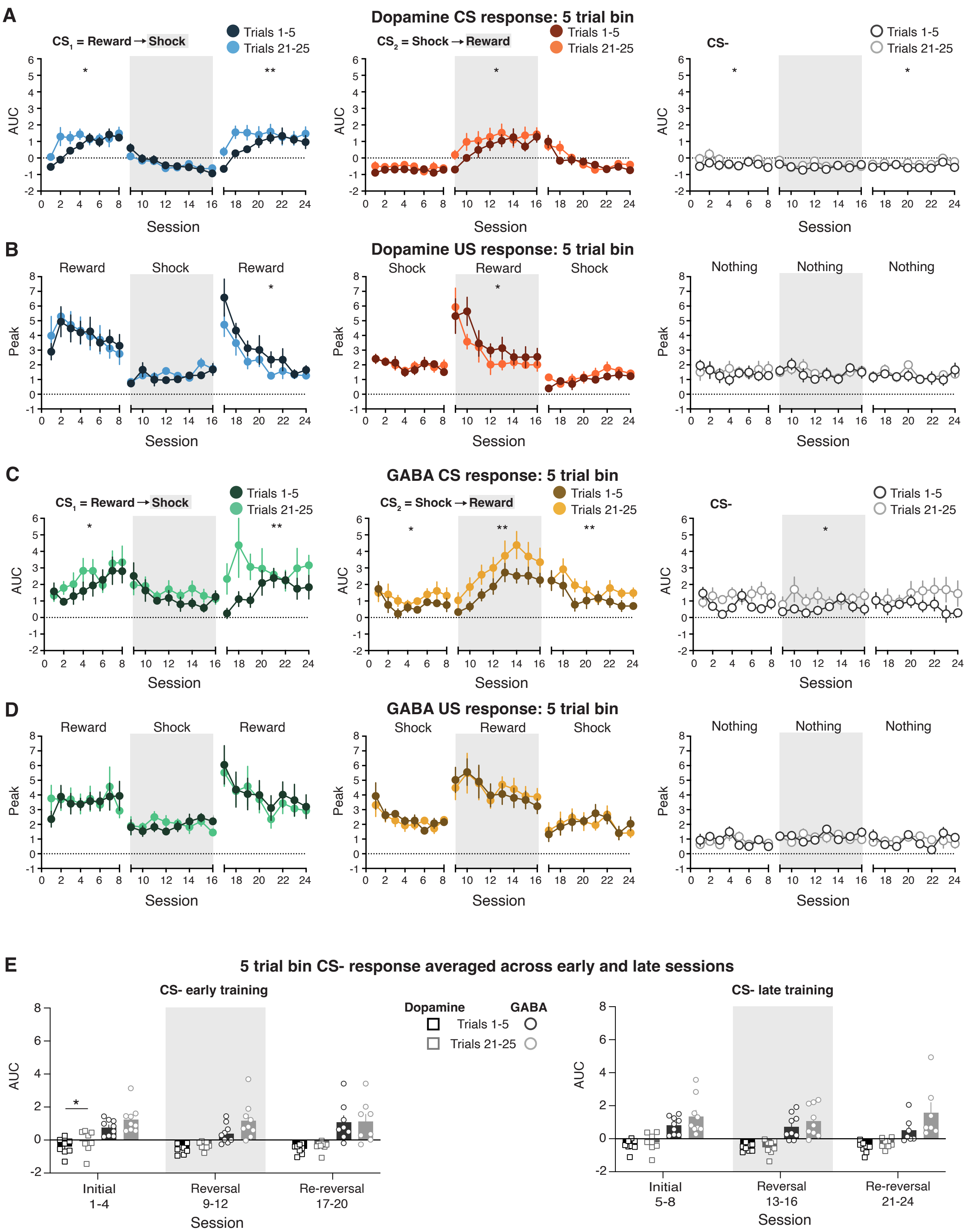


*Supplemental Figure 6*. Within-session changes in calcium activity in VTA dopamine and GABA populations across FCL. **A**, Dopamine average calcium activity during the 5 s CS in 5-trial bins across FCL. The color intensity represents the beginning of the session (Trials 1-5, dark shades) or the end of the session (Trials 21-25, light shades). **B**, Dopamine peak calcium activity in the 3 s following US delivery in 5-trial bins across FCL. **C**, GABA average calcium activity during the 5 s CS in 5-trial bins across FCL. **D**, GABA peak calcium activity in the 3 s following US delivery in 5-trial bins across FCL. **E**, Dopamine and

GABA average calcium activity during the 5 s CS-, sessions were split into 5-trial bins and bins were averaged across early (left) and late (right) sessions. The color intensity represents the beginning of the session (Trials 1-5, dark shades) or the end of the session (Trials 21-25, light shades). Data are presented as mean +/- SEM, with individual subject points in (E). *$p < 0.05$; **$p < 0.01$.

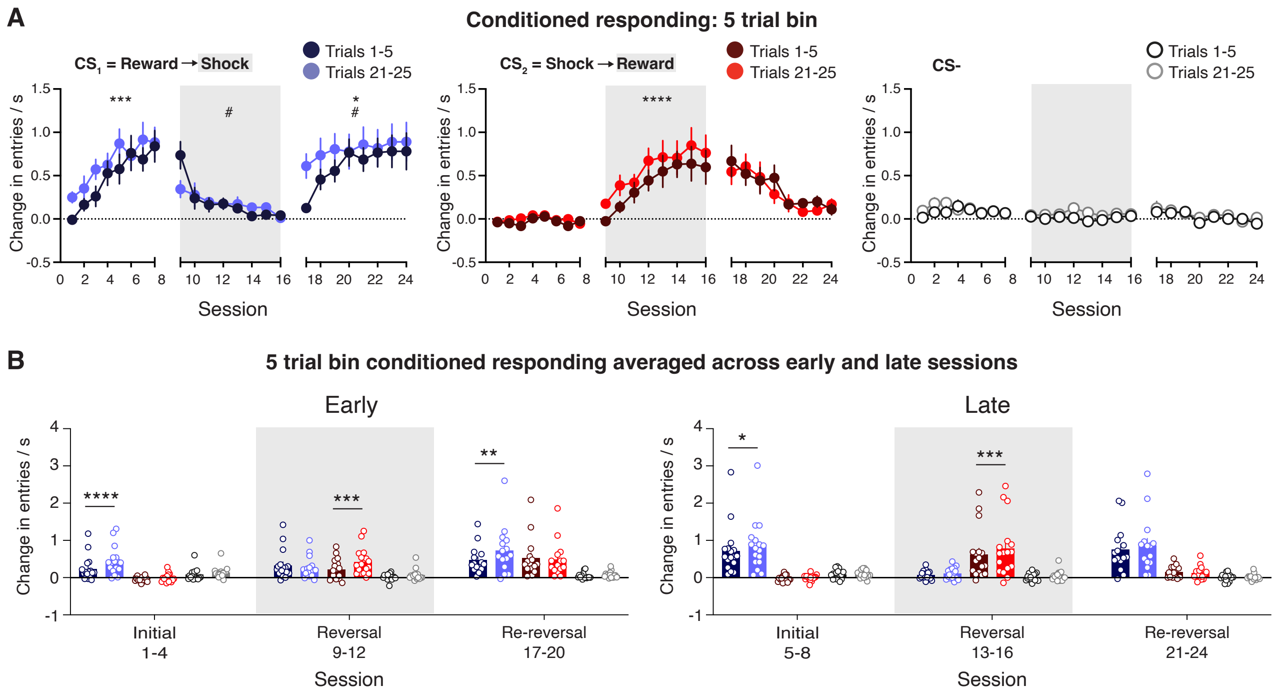


*Supplemental Figure 7*. Within-session changes in conditioned responding behavior across FCL. **A**, Average conditioned responding during the 5 s CS in 5-trial bins across FCL. The color intensity represents the beginning of the session (Trials 1-5, dark shades) or the end of the session (Trials 21-25, light shades). **B**, Within-session conditioned responding during the 5 s CS period in early (left) and late (right) sessions. Sessions were split into 5-trial bins, and bins were averaged across sessions. The color intensity represents the beginning of the session (Trials 1-5, dark shades) or the end of the session (Trials 21-25, light shades). Data are presented as mean +/- SEM, with individual subject points in (B). *$p < 0.05$; **$p < 0.01$; *** $p < 0.001$; **** $p < 0.0001$.

| Table 1: Figure 1 – FCL Behavioral Responding | | | |
|---|---|---|---|
| **Panel B: Conditioned Responding** | | | |
| Sessions 1-8<br>Two-way mixed-effects model | Session<br>$F_{(2.60, 41.63)}$ = 9.19, **$p$ = 0.0002** | CS<br>$F_{(1.14, 18.16)}$ = 22.71, **$p$ < 0.0001** | Interaction<br>$F_{(2.60, 41.66)}$ = 11.81, **$p$ < 0.0001** |
| Sessions 9-16<br>Two-way mixed-effects model | Session<br>$F_{(3.08, 49.28)}$ = 1.93, $p$ = 0.14 | CS<br>$F_{(1.06, 16.98)}$ = 12.22, **$p$ = 0.002** | Interaction<br>$F_{(1.75, 26.90)}$ = 11.97, **$p$ = 0.0003** |
| Sessions 17-24<br>Two-way mixed-effects model | Session<br>$F_{(2.76, 38.60)}$ = 1.74, $p$ = 0.18 | CS<br>$F_{(1.13, 15.79)}$ = 18.99, **$p$ = 0.0004** | Interaction<br>$F_{(1.73, 23.05)}$ = 9.15, **$p$ = 0.002** |
| Panel B: Conditioned Responding analyzed by Sex | | | |
| Sessions 1-8: Appetitive CS<br>Two-way mixed-effects model | Session<br>$F_{(7, 105)}$ = 12.81, **$p$ < 0.0001** | Sex<br>$F_{(1, 15)}$ = 2.98, $p$ = 0.11 | Interaction<br>$F_{(7, 105)}$ = 0.91, $p$ = 0.50 |
| Sessions 9-16: Appetitive CS<br>Two-way mixed-effects model | Session<br>$F_{(1.74, 25.42)}$ = 8.77, **$p$ < 0.002** | Sex<br>$F_{(1, 15)}$ = 0.76, $p$ = 0.40 | Interaction<br>$F_{(7, 102)}$ = 0.31, $p$ = 0.95 |
| **Panel C: Probability of approach** | | | |
| Sessions 1-8<br>Two-way mixed-effects model | Session<br>$F_{(3.02, 48.31)}$ = 5.75, **$p$ = 0.0019** | CS<br>$F_{(1.61, 25.69)}$ = 51.20, **$p$ < 0.0001** | Interaction<br>$F_{(5.33, 85.35)}$ = 10.25, **$p$ < 0.0001** |
| Sessions 9-16<br>Two-way mixed-effects model | Session<br>$F_{(2.66, 42.50)}$ = 0.88, $p$ = 0.45 | CS<br>$F_{(1.99, 31.87)}$ = 34.49, **$p$ < 0.0001** | Interaction<br>$F_{(4.94, 75.79)}$ = 13.61, **$p$ < 0.0001** |
| Sessions 17-24<br>Two-way mixed-effects model | Session<br>$F_{(3.21, 44.95)}$ = 4.56, **$p$ = 0.006** | CS<br>$F_{(1.89, 26.44)}$ = 40.29, **$p$ < 0.0001** | Interaction<br>$F_{(4.24, 56.67)}$ = 7.14, **$p$ < 0.0001** |
| **Panel D: Latency to approach** | | | |
| Sessions 1-8<br>Two-way mixed-effects model | Session<br>$F_{(4.82, 77.14)}$ = 1.26, $p$ = 0.29 | CS<br>$F_{(1.81, 28.98)}$ = 0.54, $p$ = 0.57 | Interaction<br>$F_{(6.85, 104.7)}$ = 2.26, **$p$ = 0.04** |
| Sessions 9-16<br>Two-way mixed-effects model | Session<br>$F_{(3.71, 59.31)}$ = 2.96, **$p$ = 0.03** | CS<br>$F_{(1.88, 30.07)}$ = 0.56, $p$ = 0.57 | Interaction<br>$F_{(6.46, 96.46)}$ = 1.47, $p$ = 0.19 |
| Sessions 17-24<br>Two-way mixed-effects model | Session<br>$F_{(3.96, 55.38)}$ = 0.79, $p$ = 0.53 | CS<br>$F_{(1.95, 27.32)}$ = 4.95, **$p$ = 0.02** | Interaction<br>$F_{(4.54, 60.65)}$ = 0.32, $p$ = 0.09 |
| **Panel E: Post-outcome head entries** | | | |
| Sessions 1-8<br>Two-way mixed-effects model | Session<br>$F_{(3.29, 52.67)}$ = 2.48, $p$ = 0.0661 | CS<br>$F_{(1.36, 21.79)}$ = 81.44, **$p$ < 0.0001** | Interaction<br>$F_{(5.21, 83.38)}$ = 7.03, **$p$ < 0.0001** |
| Sessions 9-16<br>Two-way mixed-effects model | Session<br>$F_{(2.82, 45.18)}$ = 1.49, $p$ = 0.23 | CS<br>$F_{(1.16, 18.54)}$ = 61.81, **$p$ < 0.0001** | Interaction<br>$F_{(4.39, 67.40)}$ = 1.60, $p$ **=** 0.18 |
| Sessions 17-24<br>Two-way mixed-effects model | Session<br>$F_{(3.43, 48.08)}$ = 5.54, **$p$ = 0.0016** | CS<br>$F_{(1.66, 23.24)}$ = 41.51, **$p$ < 0.0001** | Interaction<br>$F_{(4.29, 57.35)}$ = 5.40, **$p$ = 0.0007** |

| Table 2: Figure 4 – Quantified Signaling | | | |
|---|---|---|---|
| **Panel A: Dopamine full CS response** | | | |
| Sessions 1-8<br>Two-way mixed-effects model | Session<br>$F_{(2.73, 19.09)} = 3.70$, **$p = 0.03$** | CS<br>$F_{(1.82, 12.79)} = 43.07$, **$p < 0.0001$** | Interaction<br>$F_{(3.28, 21.52)} = 5.42$, **$p = 0.005$** |
| Sessions 9-16<br>Two-way mixed-effects model | Session<br>$F_{(2.67, 18.69)} = 0.78$, $p = 0.51$ | CS<br>$F_{(1.19, 8.32)} = 21.34$, **$p = 0.001$** | Interaction<br>$F_{(2.83, 17.96)} = 10.30$, **$p = 0.0004$** |
| Sessions 17-24<br>Two-way mixed-effects model | Session<br>$F_{(2.02, 14.15)} = 2.40$, $p = 0.13$ | CS<br>$F_{(1.33, 9.33)} = 22.81$, **$p = 0.0005$** | Interaction<br>$F_{(2.15, 11.81)} = 10.61$, **$p = 0.002$** |
| **Panel B: GABA full CS response** | | | |
| Sessions 1-8<br>Two-way mixed-effects model | Session<br>$F_{(1.97, 15.77)} = 1.98$, $p = 0.17$ | CS<br>$F_{(1.30, 10.37)} = 3.77$, $p = 0.07$ | Interaction<br>$F_{(2.68, 20.29)} = 2.15$, $p = 0.13$ |
| Sessions 9-16<br>Two-way mixed-effects model | Session<br>$F_{(2.73, 21.83)} = 3.23$, **$p < 0.05$** | CS<br>$F_{(1.44, 11.50)} = 6.00$, **$p = 0.02$** | Interaction<br>$F_{(2.34, 16.71)} = 7.82$, **$p = 0.003$** |
| Sessions 17-24<br>Two-way mixed-effects model | Session<br>$F_{(3.18, 22.27)} = 1.33$, $p = 0.29$ | CS<br>$F_{(1.03, 7.23)} = 4.32$, $p = 0.07$ | Interaction<br>$F_{(2.60, 13.20)} = 4.31$, **$p = 0.03$** |
| **Panel C: Dopamine early vs late CS** | | | |
| Sessions 1-8: $CS_1$<br>Two-way mixed-effects model | Session<br>$F_{(2.40, 16.77)} = 6.42$, **$p = 0.006$** | Early v late CS<br>$F_{(1, 7)} = 21.87$, **$p = 0.002$** | Interaction<br>$F_{(2.45, 15.78)} = 5.74$, **$p = 0.01$** |
| Sessions 1-8: $CS_2$<br>Two-way mixed-effects model | Session<br>$F_{(3.33, 23.30)} = 1.05$, $p = 0.39$ | Early v late CS<br>$F_{(1, 7)} = 19.91$, **$p = 0.003$** | Interaction<br>$F_{(3.81, 24.51)} = 0.19$, $p = 0.93$ |
| Sessions 1-8: CS-<br>Two-way mixed-effects model | Session<br>$F_{(3.08, 21.52)} = 1.75$, $p = 0.19$ | Early v late<br>$F_{(1, 7)} = 10.04$, **$p = 0.02$** | Interaction<br>$F_{(1.79, 11.54)} = 0.61$, $p = 0.55$ |
| Sessions 9-16: $CS_1$<br>Two-way mixed-effects model | Session<br>$F_{(1.68, 11.73)} = 8.14$, **$p = 0.008$** | Early v late CS<br>$F_{(1, 7)} = 13.36$, **$p = 0.008$** | Interaction<br>$F_{(2.49, 15.29)} = 5.00$, **$p = 0.02$** |
| Sessions 9-16: $CS_2$<br>Two-way mixed-effects model | Session<br>$F_{(2.47, 17.31)} = 8.08$, **$p = 0.002$** | Early v late CS<br>$F_{(1, 7)} = 8.56$, **$p = 0.02$** | Interaction<br>$F_{(2.53, 15.53)} = 3.63$, **$p = 0.04$** |
| Sessions 9-16: CS-<br>Two-way mixed-effects model | Session<br>$F_{(2.90, 20.30)} = 1.45$, $p = 0.26$ | Early v late CS<br>$F_{(1, 7)} = 0.00002$, $p = 0.99$ | Interaction<br>$F_{(2.63, 16.13)} = 0.90$, $p = 0.45$ |
| Sessions 17-24: $CS_1$<br>Two-way mixed-effects model | Session<br>$F_{(1.35, 9.48)} = 6.76$, **$p = 0.02$** | Early v late CS<br>$F_{(1, 7)} = 4.19$, $p = 0.08$ | Interaction<br>$F_{(1.69, 8.46)} = 7.82$, **$p = 0.01$** |
| Sessions 17-24: $CS_2$<br>Two-way mixed-effects model | Session<br>$F_{(1.31, 9.19)} = 12.21$, **$p = 0.005$** | Early v late CS<br>$F_{(1, 7)} = 9.07$, **$p = 0.02$** | Interaction<br>$F_{(1.98, 9.89)} = 2.66$, $p = 0.12$ |
| Sessions 17-24: CS-<br>Two-way mixed-effects model | Session<br>$F_{(1.87, 13.07)} = 1.47$, $p = 0.26$ | Early v late CS<br>$F_{(1, 7)} = 5.33$, $p = 0.05$ | Interaction<br>$F_{(3.36, 16.79)} = 0.56$, $p = 0.67$ |
| **Panel D: GABA early vs late CS** | | | |
| Sessions 1-8: $CS_1$<br>Two-way mixed-effects model | Session<br>$F_{(1.95, 15.63)} = 2.81$, $p = 0.09$ | Early v late CS<br>$F_{(1, 8)} = 7.63$, **$p = 0.02$** | Interaction<br>$F_{(1.85, 13.81)} = 0.66$, $p = 0.52$ |
| Sessions 1-8: $CS_2$<br>Two-way mixed-effects model | Session<br>$F_{(1.94, 15.50)} = 0.89$, $p = 0.43$ | Early v late CS<br>$F_{(1, 8)} = 9.71$, **$p = 0.01$** | Interaction<br>$F_{(2.45, 18.17)} = 1.24$, $p = 0.32$ |
| Sessions 1-8: CS-<br>Two-way mixed-effects model | Session<br>$F_{(2.00, 16.04)} = 0.62$, $p = 0.55$ | Early v late CS<br>$F_{(1, 8)} = 14.35$, **$p = 0.005$** | Interaction<br>$F_{(2.65, 19.70)} = 0.34$, $p = 0.77$ |
| Sessions 9-16: $CS_1$<br>Two-way mixed-effects model | Session<br>$F_{(1.94, 15.51)} = 2.15$, $p = 0.15$ | Early v late CS<br>$F_{(1, 8)} = 8.43$, **$p = 0.02$** | Interaction<br>$F_{(1.42, 9.70)} = 1.60$, $p = 0.25$ |
| Sessions 9-16: $CS_2$<br>Two-way mixed-effects model | Session<br>$F_{(3.22, 24.85)} = 9.56$, **$p = 0.0002$** | Early v late CS<br>$F_{(1, 8)} = 0.66$, $p = 0.44$ | Interaction<br>$F_{(2.33, 15.97)} = 1.12$, $p = 0.36$ |
| Sessions 9-16: CS-<br>Two-way mixed-effects model | Session<br>$F_{(2.14, 17.10)} = 1.07$, $p = 0.37$ | Early v late CS<br>$F_{(1, 8)} = 4.41$, $p = 0.07$ | Interaction<br>$F_{(2.27, 15.58)} = 1.21$, $p = 0.33$ |
| Sessions 17-24: $CS_1$<br>Two-way mixed-effects model | Session<br>$F_{(2.84, 19.89)} = 2.76$, $p = 0.07$ | Early v late CS<br>$F_{(1, 7)} = 0.77$, $p = 0.41$ | Interaction<br>$F_{(2.11, 9.36)} = 0.78$, $p = 0.49$ |
| Sessions 17-24: $CS_2$<br>Two-way mixed-effects model | Session<br>$F_{(2.52, 17.67)} = 3.82$, **$p = 0.03$** | Early v late CS<br>$F_{(1, 7)} = 4.08$, $p = 0.08$ | Interaction<br>$F_{(2.34, 10.38)} = 0.81$, $p = 0.49$ |
| Sessions 17-24: CS-<br>Two-way mixed-effects model | Session<br>$F_{(1.79, 12.50)} = 1.28$, $p = 0.31$ | Early v late CS<br>$F_{(1, 7)} = 5.06$, $p = 0.06$ | Interaction<br>$F_{(2.62, 11.61)} = 1.31$, $p = 0.31$ |

| Table 3: Figure 5 – Quantified Signaling – 5 trial bins | | | |
|---|---|---|---|
| **Panel C: Dopamine 5 trial bin CS response averaged across early phases of training** | | | |
| $CS_1$ = Reward -> Shock<br>Two-way mixed-effects model | Training phase<br>$F_{(1.53, 10.76)} = 6.01$, $\boldsymbol{p = 0.02}$ | Trials<br>$F_{(1, 7)} = 28.22$, $\boldsymbol{p = 0.001}$ | Interaction<br>$F_{(1.96, 13.69)} = 9.95$, $\boldsymbol{p = 0.002}$ |
| $CS_2$ = Shock -> Reward<br>Two-way mixed-effects model | Training phase<br>$F_{(1.37, 9.59)} = 10.37$, $\boldsymbol{p = 0.007}$ | Trials<br>$F_{(1, 7)} = 7.75$, $\boldsymbol{p = 0.03}$ | Interaction<br>$F_{(1.35, 9.43)} = 4.78$, $\boldsymbol{p < 0.05}$ |
| **Panel D: Dopamine 5 trial bin CS response averaged across late phases of training** | | | |
| $CS_1$ = Reward -> Shock<br>Two-way mixed-effects model | Training phase<br>$F_{(1.83, 12.83)} = 30.40$, $\boldsymbol{p < 0.0001}$ | Trials<br>$F_{(1, 7)} = 0.96$, $p = 0.36$ | Interaction<br>$F_{(1.28, 8.93)} = 0.30$, $p = 0.65$ |
| $CS_2$ = Shock -> Reward<br>Two-way mixed-effects model | Training phase<br>$F_{(1.07, 7.52)} = 20.69$, $\boldsymbol{p = 0.002}$ | Trials<br>$F_{(1, 7)} = 3.30$, $p = 0.11$ | Interaction<br>$F_{(1.47, 10.29)} = 1.30$, $p = 0.30$ |
| **Panel E: GABA 5 trial bin CS response averaged across early phases of training** | | | |
| $CS_1$ = Reward -> Shock<br>Two-way mixed-effects model | Training phase<br>$F_{(1.38, 11.07)} = 3.46$, $p = 0.08$ | Trials<br>$F_{(1, 8)} = 20.33$, $\boldsymbol{p = 0.002}$ | Interaction<br>$F_{(1.33, 9.28)} = 8.22$, $\boldsymbol{p = 0.01}$ |
| $CS_2$ = Shock -> Reward<br>Two-way mixed-effects model | Training phase<br>$F_{(1.65, 13.19)} = 2.28$, $p = 0.15$ | Trials<br>$F_{(1, 8)} = 15.86$, $\boldsymbol{p = 0.004}$ | Interaction<br>$F_{(1.62, 11.31)} = 1.86$, $p = 0.20$ |
| **Panel F: GABA 5 trial bin CS response averaged across late phases of training** | | | |
| $CS_1$ = Reward -> Shock<br>Two-way mixed-effects model | Training phase<br>$F_{(1.94, 15.53)} = 7.45$, $\boldsymbol{p = 0.006}$ | Trials<br>$F_{(1, 8)} = 14.37$, $\boldsymbol{p = 0.005}$ | Interaction<br>$F_{(1.61, 9.64)} = 0.50$, $p = 0.58$ |
| $CS_2$ = Shock -> Reward<br>Two-way mixed-effects model | Training phase<br>$F_{(1.35, 10.76)} = 13.76$, $\boldsymbol{p = 0.002}$ | Trials<br>$F_{(1, 8)} = 29.74$, $\boldsymbol{p = 0.0006}$ | Interaction<br>$F_{(1.56, 9.36)} = 2.50$, $p = 0.14$ |

| Supplemental Table 1: Supplemental Figure 1 – FCL Behavioral Responding | | | |
|---|---|---|---|
| **Panel A: Conditioned Responding to Appetitive association** | | | |
| Two-way mixed-effects model | Session<br>$F_{(1.54, 24.69)}$ = 14.31, ***p* = 0.0002** | Training phase<br>$F_{(1.22, 19.49)}$ = 2.98, *p* = 0.09 | Interaction<br>$F_{(4.23, 61.09)}$ = 1.19, *p* = 0.33 |
| **Panel B: Conditioned Responding analyzed by Genotype** | | | |
| Sessions 1-8: $CS_1$<br>Two-way mixed-effects model | Session<br>$F_{(2.05, 30.69)}$ = 12.03, ***p* = 0.0001** | Genotype<br>$F_{(1, 15)}$ = 0.09, *p* = 0.77 | Interaction<br>$F_{(7, 105)}$ = 0.47, *p* = 0.85 |
| Sessions 1-8: $CS_2$<br>Two-way mixed-effects model | Session<br>$F_{(3.00, 44.97)}$ = 2.58, *p* = 0.07 | Genotype<br>$F_{(1, 15)}$ = 0.42, *p* = 0.53 | Interaction<br>$F_{(7, 105)}$ = 1.12, *p* = 0.36 |
| Sessions 1-8: CS-<br>Two-way mixed-effects model | Session<br>$F_{(3.02, 45.23)}$ = 1.52, *p* = 0.22 | Genotype<br>$F_{(1, 15)}$ = 1.06, *p* = 0.32 | Interaction<br>$F_{(7, 105)}$ = 1.19, *p* = 0.31 |
| Sessions 9-16: $CS_1$<br>Two-way mixed-effects model | Session<br>$F_{(1.69, 24.56)}$ = 8.46, ***p* = 0.003** | Genotype<br>$F_{(1, 15)}$ = 0.15, *p* = 0.70 | Interaction<br>$F_{(7, 102)}$ = 0.59, *p* = 0.76 |
| Sessions 9-16: $CS_2$<br>Two-way mixed-effects model | Session<br>$F_{(2.12, 30.88)}$ = 9.75, ***p* = 0.0004** | Genotype<br>$F_{(1, 15)}$ = 1.62, *p* = 0.22 | Interaction<br>$F_{(7, 102)}$ = 1.95, *p* = 0.07 |
| Sessions 9-16: CS-<br>Two-way mixed-effects model | Session<br>$F_{(2.78, 40.53)}$ = 0.63, *p* = 0.59 | Genotype<br>$F_{(1, 15)}$ = 0.15, *p* = 0.70 | Interaction<br>$F_{(7, 102)}$ = 1.01, *p* = 0.43 |
| Sessions 17-24: $CS_1$<br>Two-way mixed-effects model | Session<br>$F_{(3.06, 38.52)}$ = 7.13, ***p* = 0.0006** | Genotype<br>$F_{(1, 13)}$ = 3.05, *p* = 0.10 | Interaction<br>$F_{(7, 88)}$ = 4.60, ***p* = 0.002** |
| Sessions 17-24: $CS_2$<br>Two-way mixed-effects model | Session<br>$F_{(1.69, 21.27)}$ = 7.75, ***p* = 0.004** | Genotype<br>$F_{(1, 13)}$ = 0.45, *p* = 0.51 | Interaction<br>$F_{(7, 88)}$ = 1.78, *p* = 0.10 |
| Sessions 17-24: CS-<br>Two-way mixed-effects model | Session<br>$F_{(3.54, 44.52)}$ = 2.98, ***p* = 0.03** | Genotype<br>$F_{(1, 13)}$ = 0.12, *p* = 0.73 | Interaction<br>$F_{(7, 88)}$ = 0.75, *p* = 0.63 |
| **Panel C: Conditioned Responding analyzed by Sex** | | | |
| Sessions 1-8: $CS_1$<br>Two-way mixed-effects model | Session<br>$F_{(2.07, 31.04)}$ = 12.81, ***p* < 0.0001** | Sex<br>$F_{(1, 15)}$ = 2.98, *p* = 0.11 | Interaction<br>$F_{(7, 105)}$ = 0.91, *p* = 0.50 |
| Sessions 1-8: $CS_2$<br>Two-way mixed-effects model | Session<br>$F_{(3.02, 45.27)}$ = 2.12, *p* = 0.11 | Sex<br>$F_{(1, 15)}$ = 0.37, *p* = 0.55 | Interaction<br>$F_{(7, 105)}$ = 1.46, *p* = 0.19 |
| Sessions 1-8: CS-<br>Two-way mixed-effects model | Session<br>$F_{(7, 105)}$ = 1.44, *p* = 0.20 | Sex<br>$F_{(1, 15)}$ = 0.43, *p* = 0.52 | Interaction<br>$F_{(7, 105)}$ = 0.72, *p* = 0.66 |
| Sessions 9-16: $CS_1$<br>Two-way mixed-effects model | Session<br>$F_{(1.74, 25.30)}$ = 8.54, ***p* = 0.002** | Sex<br>$F_{(1, 15)}$ = 2.77, *p* = 0.12 | Interaction<br>$F_{(7, 102)}$ = 0.23, *p* = 0.97 |
| Sessions 9-16: $CS_2$<br>Two-way mixed-effects model | Session<br>$F_{(1.74, 25.42)}$ = 8.77, ***p* = 0.002** | Sex<br>$F_{(1, 15)}$ = 0.76, *p* = 0.40 | Interaction<br>$F_{(7, 102)}$ = 0.31, *p* = 0.95 |
| Sessions 9-16: CS-<br>Two-way mixed-effects model | Session<br>$F_{(7, 102)}$ = 0.50, *p* = 0.83 | Sex<br>$F_{(1, 15)}$ = 0.84, *p* = 0.37 | Interaction<br>$F_{(7, 102)}$ = 0.56, *p* = 0.79 |
| Sessions 17-24: $CS_1$<br>Two-way mixed-effects model | Session<br>$F_{(2.27, 28.51)}$ = 5.01, ***p* = 0.01** | Sex<br>$F_{(1, 13)}$ = 0.19, *p* = 0.67 | Interaction<br>$F_{(7, 88)}$ = 0.49, *p* = 0.84 |
| Sessions 17-24: $CS_2$<br>Two-way mixed-effects model | Session<br>$F_{(1.53, 19.26)}$ = 7.46, ***p* = 0.007** | Sex<br>$F_{(1, 13)}$ = 0.06, *p* = 0.81 | Interaction<br>$F_{(7, 88)}$ = 0.66, *p* = 0.70 |
| Sessions 17-24: CS-<br>Two-way mixed-effects model | Session<br>$F_{(7, 88)}$ = 3.05, ***p* = 0.006** | Sex<br>$F_{(1, 13)}$ = 2.38, *p* = 0.15 | Interaction<br>$F_{(7, 88)}$ = 0.47, *p* = 0.85 |

| Supplemental Table 2: Supplemental Figure 5 – TH and GAD signaling | | | |
|---|---|---|---|
| **Panel A: Full CS response by Genotype** | | | |
| Sessions 1-8: $CS_1$<br>Two-way mixed-effects model | Session<br>$F_{(7, 101)} = 5.49$, $\boldsymbol{p < 0.0001}$ | Genotype<br>$F_{(1, 15)} = 5.86$, $\boldsymbol{p = 0.03}$ | Interaction<br>$F_{(7, 101)} = 0.71$, $p = 0.66$ |
| Sessions 1-8: $CS_2$<br>Two-way mixed-effects model | Session<br>$F_{(2.13, 30.74)} = 0.64$, $p = 0.55$ | Genotype<br>$F_{(1, 15)} = 33.02$, $\boldsymbol{p < 0.0001}$ | Interaction<br>$F_{(7, 101)} = 1.03$, $p = 0.41$ |
| Sessions 1-8: CS-<br>Two-way mixed-effects model | Session<br>$F_{(2.47, 25.70)} = 0.21$, $p = 0.85$ | Genotype<br>$F_{(1, 15)} = 21.36$, $\boldsymbol{p = 0.0003}$ | Interaction<br>$F_{(7, 101)} = 1.47$, $p = 0.19$ |
| Sessions 9-16: $CS_1$<br>Two-way mixed-effects model | Session<br>$F_{(7, 98)} = 5.54$, $\boldsymbol{p < 0.0001}$ | Genotype<br>$F_{(1, 15)} = 30.97$, $\boldsymbol{p < 0.0001}$ | Interaction<br>$F_{(7, 98)} = 0.50$, $p = 0.83$ |
| Sessions 9-16: $CS_2$<br>Two-way mixed-effects model | Session<br>$F_{(3.66, 51.23)} = 15.78$, $\boldsymbol{p < 0.0001}$ | Genotype<br>$F_{(1, 15)} = 7.20$, $\boldsymbol{p = 0.02}$ | Interaction<br>$F_{(7, 98)} = 1.39$, $p = 0.22$ |
| Sessions 9-16: CS-<br>Two-way mixed-effects model | Session<br>$F_{(2.89, 40.49)} = 1.05$, $p = 0.38$ | Genotype<br>$F_{(1, 15)} = 17.51$, $\boldsymbol{p = 0.0008}$ | Interaction<br>$F_{(7, 98)} = 1.14$, $p = 0.35$ |
| Sessions 17-24: $CS_1$<br>Two-way mixed-effects model | Session<br>$F_{(7, 82)} = 6.63$, $\boldsymbol{p < 0.0001}$ | Genotype<br>$F_{(1, 14)} = 5.93$, $\boldsymbol{p = 0.03}$ | Interaction<br>$F_{(7, 82)} = 0.78$, $p = 0.60$ |
| Sessions 17-24: $CS_2$<br>Two-way mixed-effects model | Session<br>$F_{(2.32, 27.16)} = 11.88$, $\boldsymbol{p = 0.0001}$ | Genotype<br>$F_{(1, 14)} = 33.37$, $\boldsymbol{p < 0.0001}$ | Interaction<br>$F_{(7, 82)} = 0.89$, $p = 0.52$ |
| Sessions 17-24: CS-<br>Two-way mixed-effects model | Session<br>$F_{(2.05, 23.96)} = 1.66$, $p = 0.21$ | Genotype<br>$F_{(1, 14)} = 12.85$, $\boldsymbol{p = 0.003}$ | Interaction<br>$F_{(7, 82)} = 0.97$, $p = 0.46$ |
| **Panel B: Dopamine US peak** | | | |
| Sessions 1-8<br>Two-way mixed-effects model | Session<br>$F_{(1.49, 10.41)} = 2.32$, $p = 0.15$ | US<br>$F_{(1.15, 8.03)} = 19.48$, $\boldsymbol{p = 0.002}$ | Interaction<br>$F_{(2.14, 14.09)} = 2.79$, $p = 0.09$ |
| Sessions 9-16<br>Two-way mixed-effects model | Session<br>$F_{(2.41, 16.90)} = 7.99$, $\boldsymbol{p = 0.003}$ | US<br>$F_{(1.21, 8.50)} = 10.89$, $\boldsymbol{p = 0.008}$ | Interaction<br>$F_{(2.42, 15.36)} = 13.14$, $\boldsymbol{p = 0.0003}$ |
| Sessions 17-24<br>Two-way mixed-effects model | Session<br>$F_{(1.79, 12.50)} = 9.01$, $\boldsymbol{p = 0.005}$ | US<br>$F_{(1.20, 8.41)} = 11.56$, $\boldsymbol{p = 0.007}$ | Interaction<br>$F_{(1.85, 10.16)} = 14.78$, $\boldsymbol{p = 0.001}$ |
| **Panel C: GABA US peak** | | | |
| Sessions 1-8<br>Two-way mixed-effects model | Session<br>$F_{(2.62, 20.95)} = 0.91$, $p = 0.44$ | US<br>$F_{(1.46, 11.66)} = 21.67$, $\boldsymbol{p = 0.0003}$ | Interaction<br>$F_{(3.30, 24.97)} = 2.37$, $p = 0.09$ |
| Sessions 9-16<br>Two-way mixed-effects model | Session<br>$F_{(2.84, 22.71)} = 2.41$, $p = 0.09$ | US<br>$F_{(1.21, 9.66)} = 24.46$, $\boldsymbol{p = 0.0004}$ | Interaction<br>$F_{(4.49, 32.09)} = 2.80$, $\boldsymbol{p = 0.04}$ |
| Sessions 17-24<br>Two-way mixed-effects model | Session<br>$F_{(2.21, 15.47)} = 3.30$, $p = 0.06$ | US<br>$F_{(1.72, 12.04)} = 12.64$, $\boldsymbol{p = 0.002}$ | Interaction<br>$F_{(2.07, 10.50)} = 4.87$, $\boldsymbol{p = 0.03}$ |
| **Panel D: Dopamine full US response** | | | |
| Sessions 1-8<br>Two-way mixed-effects model | Session<br>$F_{(1.78, 12.49)} = 4.04$, $\boldsymbol{p < 0.05}$ | US<br>$F_{(1.80, 12.60)} = 2.76$, $p = 0.11$ | Interaction<br>$F_{(3.30, 21.67)} = 2.14$, $p = 0.12$ |
| Sessions 9-16<br>Two-way mixed-effects model | Session<br>$F_{(2.48, 17.38)} = 3.00$, $p = 0.07$ | US<br>$F_{(1.74, 12.16)} = 3.35$, $p = 0.07$ | Interaction<br>$F_{(2.90, 18.44)} = 15.60$, $\boldsymbol{p < 0.0001}$ |
| Sessions 17-24<br>Two-way mixed-effects model | Session<br>$F_{(2.30, 16.10)} = 2.54$, $p = 0.10$ | US<br>$F_{(1.87, 13.08)} = 2.04$, $p = 0.17$ | Interaction<br>$F_{(2.58, 14.17)} = 13.86$, $\boldsymbol{p = 0.0002}$ |
| **Panel E: GABA full US response** | | | |
| Sessions 1-8<br>Two-way mixed-effects model | Session<br>$F_{(2.62, 20.93)} = 0.68$, $p = 0.56$ | US<br>$F_{(1.21, 9.71)} = 7.43$, $\boldsymbol{p = 0.02}$ | Interaction<br>$F_{(3.88, 29.36)} = 0.79$, $p = 0.54$ |
| Sessions 9-16<br>Two-way mixed-effects model | Session<br>$F_{(2.18, 17.42)} = 4.38$, $\boldsymbol{p = 0.03}$ | US<br>$F_{(1.39, 11.15)} = 13.95$, $\boldsymbol{p = 0.002}$ | Interaction<br>$F_{(2.62, 18.72)} = 7.21$, $\boldsymbol{p = 0.003}$ |
| Sessions 17-24<br>Two-way mixed-effects model | Session<br>$F_{(2.27, 15.90)} = 2.19$, $p = 0.14$ | US<br>$F_{(1.78, 12.48)} = 5.25$, $\boldsymbol{p = 0.02}$ | Interaction<br>$F_{(1.95, 9.87)} = 4.54$, $\boldsymbol{p = 0.04}$ |

| Supplemental Table 3: Supplemental Figure 6 – TH and GAD Signaling – 5 trial bins | | | |
|---|---|---|---|
| **Panel A: Dopamine CS response (AUC) – 5 trial bin** | | | |
| **Left: $CS_1$ = Reward -> Shock** | | | |
| Sessions 1-8<br>Two-way mixed-effects model | Session<br>$F_{(2.29, 16.01)} = 5.31$, **$p = 0.01$** | Trial<br>$F_{(1, 7)} = 8.53$, **$p = 0.02$** | Interaction<br>$F_{(3.34, 21.49)} = 2.63$, $p = 0.07$ |
| Sessions 9-16<br>Two-way mixed-effects model | Session<br>$F_{(1.82, 12.71)} = 8.43$, **$p = 0.005$** | Trial<br>$F_{(1, 7)} = 0.16$, $p = 0.70$ | Interaction<br>$F_{(3.06, 18.38)} = 1.72$, $p = 0.20$ |
| Sessions 17-24<br>Two-way mixed-effects model | Session<br>$F_{(2.35, 16.48)} = 8.42$, **$p = 0.002$** | Trial<br>$F_{(1, 7)} = 12.28$, **$p = 0.009$** | Interaction<br>$F_{(2.29, 11.79)} = 1.92$, $p = 0.19$ |
| **Middle: $CS_2$ = Shock -> Reward** | | | |
| Sessions 1-8<br>Two-way mixed-effects model | Session<br>$F_{(2.28, 19.44)} = 0.39$, $p = 0.75$ | Trial<br>$F_{(1, 7)} = 3.70$, $p = 0.10$ | Interaction<br>$F_{(3.37, 21.64)} = 0.98$, $p = 0.43$ |
| Sessions 9-16<br>Two-way mixed-effects model | Session<br>$F_{(2.10, 14.67)} = 6.55$, **$p = 0.009$** | Trial<br>$F_{(1, 7)} = 6.90$, **$p = 0.03$** | Interaction<br>$F_{(2.32, 13.94)} = 1.84$, $p = 0.19$ |
| Sessions 17-24<br>Two-way mixed-effects model | Session<br>$F_{(2.29, 16.00)} = 0.33$, **$p = 0.0003$** | Trial<br>$F_{(1, 7)} = 0.49$, $p = 0.51$ | Interaction<br>$F_{(2.68, 14.16)} = 3.95$, **$p = 0.03$** |
| **Right: CS-** | | | |
| Sessions 1-8<br>Two-way mixed-effects model | Session<br>$F_{(3.32, 23.27)} = 2.29$, $p = 0.10$ | Trial<br>$F_{(1, 7)} = 10.62$, **$p = 0.01$** | Interaction<br>$F_{(3.67, 23.57)} = 1.49$, $p = 0.24$ |
| Sessions 9-16<br>Two-way mixed-effects model | Session<br>$F_{(2.86, 20.04)} = 1.47$, $p = 0.25$ | Trial<br>$F_{(1, 7)} = 1.47$, $p = 0.26$ | Interaction<br>$F_{(3.82, 22.92)} = 0.64$, $p = 0.63$ |
| Sessions 17-24<br>Two-way mixed-effects model | Session<br>$F_{(2.74, 19.20)} = 1.59$, $p = 0.23$ | Trial<br>$F_{(1, 7)} = 9.97$, **$p = 0.02$** | Interaction<br>$F_{(2.55, 13.13)} = 0.35$, $p = 0.76$ |
| **Panel B: Dopamine US response (Peak) – 5 trial bin** | | | |
| **Left: $CS_1$ = Reward -> Shock** | | | |
| Sessions 1-8<br>Two-way mixed-effects model | Session<br>$F_{(2.11, 14.75)} = 1.98$, $p = 0.17$ | Trial<br>$F_{(1, 7)} = 0.11$, $p = 0.75$ | Interaction<br>$F_{(2.65, 17.04)} = 0.72$, $p = 0.54$ |
| Sessions 9-16<br>Two-way mixed-effects model | Session<br>$F_{(3.06, 21.39)} = 2.07$, $p = 0.13$ | Trial<br>$F_{(1, 7)} = 2.09$, $p = 0.19$ | Interaction<br>$F_{(2.84, 17.05)} = 1.22$, $p = 0.33$ |
| Sessions 17-24<br>Two-way mixed-effects model | Session<br>$F_{(1.73, 12.12)} = 12.67$, **$p = 0.001$** | Trial<br>$F_{(1, 7)} = 7.60$, **$p = 0.03$** | Interaction<br>$F_{(2.21, 11.36)} = 1.00$, $p = 0.40$ |
| **Middle: $CS_2$ = Shock -> Reward** | | | |
| Sessions 1-8<br>Two-way mixed-effects model | Session<br>$F_{(3.02, 21.11)} = 2.44$, $p = 0.09$ | Trial<br>$F_{(1, 7)} = 0.32$, $p = 0.59$ | Interaction<br>$F_{(4.04, 25.95)} = 0.46$, $p = 0.77$ |
| Sessions 9-16<br>Two-way mixed-effects model | Session<br>$F_{(1.86, 13.03)} = 10.04$, **$p = 0.003$** | Trial<br>$F_{(1, 7)} = 5.93$, **$p < 0.05$** | Interaction<br>$F_{(2.84, 17.53)} = 1.44$, $p = 0.26$ |
| Sessions 17-24<br>Two-way mixed-effects model | Session<br>$F_{(3.14, 21.97)} = 2.05$, $p = 0.13$ | Trial<br>$F_{(1, 7)} = 4.98$, $p = 0.06$ | Interaction<br>$F_{(3.84, 20.28)} = 0.98$, $p = 0.44$ |
| **Right: CS-** | | | |
| Sessions 1-8<br>Two-way mixed-effects model | Session<br>$F_{(2.95, 20.62)} = 0.50$, $p = 0.68$ | Trial<br>$F_{(1, 7)} = 2.37$, $p = 0.17$ | Interaction<br>$F_{(3.46, 22.27)} = 1.34$, $p = 0.29$ |
| Sessions 9-16<br>Two-way mixed-effects model | Session<br>$F_{(2.75, 19.23)} = 1.38$, $p = 0.28$ | Trial<br>$F_{(1, 7)} = 0.67$, $p = 0.44$ | Interaction<br>$F_{(3.38, 20.26)} = 1.35$, $p = 0.29$ |
| Sessions 17-24<br>Two-way mixed-effects model | Session<br>$F_{(2.65, 18.52)} = 1.94$, $p = 0.16$ | Trial<br>$F_{(1, 7)} = 0.54$, $p = 0.49$ | Interaction<br>$F_{(2.67, 13.73)} = 1.01$, $p = 0.41$ |
| **Panel C: GABA CS response (AUC) – 5 trial bin** | | | |
| **Left: CS1 = Reward -> Shock** | | | |
| Sessions 1-8<br>Two-way mixed-effects model | Session<br>$F_{(2.07, 16.59)} = 2.58$, $p = 0.10$ | Trial<br>$F_{(1, 8)} = 5.28$, **$p < 0.05$** | Interaction<br>$F_{(3.19, 23.69)} = 1.29$, $p = 0.30$ |
| Sessions 9-16<br>Two-way mixed-effects model | Session<br>$F_{(1.99, 15.91)} = 3.17$, $p = 0.07$ | Trial<br>$F_{(1, 8)} = 4.47$, $p = 0.07$ | Interaction<br>$F_{(3.07, 19.71)} = 1.18$, $p = 0.35$ |
| Sessions 17-24<br>Two-way mixed-effects model | Session<br>$F_{(2.29, 16.02)} = 0.93$, $p = 0.43$ | Trial<br>$F_{(1, 7)} = 23.26$, **$p = 0.002$** | Interaction<br>$F_{(1.92, 8.22)} = 1.56$, $p = 0.27$ |
| **Middle: CS2 = Shock -> Reward** | | | |
| Sessions 1-8<br>Two-way mixed-effects model | Session<br>$F_{(2.35, 18.76)} = 1.70$, $p = 0.21$ | Trial<br>$F_{(1, 8)} = 8.81$, **$p = 0.02$** | Interaction<br>$F_{(2.65, 19.68)} = 1.00$, $p = 0.41$ |
| Sessions 9-16 | Session | Trial | Interaction |

| | | | |
|---|---|---|---|
| Two-way mixed-effects model | $F_{(3.65, 29.20)} = 7.21$, ***p* = 0.0005** | $F_{(1, 8)} = 17.72$, ***p* = 0.003** | $F_{(2.25, 14.17)} = 0.32$, $p = 0.76$ |
| Sessions 17-24<br>Two-way mixed-effects model | Session<br>$F_{(2.34, 16.37)} = 3.85$, ***p* = 0.04** | Trial<br>$F_{(1, 7)} = 23.01$, ***p* = 0.002** | Interaction<br>$F_{(2.65, 11.75)} = 1.41$, $p = 0.29$ |
| **Right: CS-** | | | |
| Sessions 1-8<br>Two-way mixed-effects model | Session<br>$F_{(2.41, 19.25)} = 0.50$, $p = 0.65$ | Trial<br>$F_{(1, 8)} = 4.32$, $p = 0.07$ | Interaction<br>$F_{(2.65, 19.68)} = 2.79$, $p = 0.07$ |
| Sessions 9-16<br>Two-way mixed-effects model | Session<br>$F_{(2.89, 18.21)} = 0.61$, $p = 0.58$ | Trial<br>$F_{(1, 8)} = 10.85$, ***p* = 0.01** | Interaction<br>$F_{(1.98, 12.42)} = 1.12$, $p = 0.36$ |
| Sessions 17-24<br>Two-way mixed-effects model | Session<br>$F_{(2.40, 16.78)} = 0.81$, $p = 0.48$ | Trial<br>$F_{(1, 7)} = 2.40$, $p = 0.17$ | Interaction<br>$F_{(2.04, 8.74)} = 2.03$, $p = 0.19$ |
| **Panel D: GABA US response (Peak) – 5 trial bin** | | | |
| **Left: CS1 = Reward -> Shock** | | | |
| Sessions 1-8<br>Two-way mixed-effects model | Session<br>$F_{(2.63, 21.03)} = 0.51$, $p = 0.66$ | Trial<br>$F_{(1, 8)} = 0.25$, $p = 0.63$ | Interaction<br>$F_{(3.10, 23.03)} = 0.76$, $p = 0.53$ |
| Sessions 9-16<br>Two-way mixed-effects model | Session<br>$F_{(2.08, 16.67)} = 0.64$, $p = 0.54$ | Trial<br>$F_{(1, 8)} = 0.20$, $p = 0.67$ | Interaction<br>$F_{(3.22, 20.72)} = 1.39$, $p = 0.27$ |
| Sessions 17-24<br>Two-way mixed-effects model | Session<br>$F_{(1.55, 10.84)} = 4.27$, $p = 0.05$ | Trial<br>$F_{(1, 7)} = 0.28$, $p = 0.62$ | Interaction<br>$F_{(2.45, 10.52)} = 0.22$, $p = 0.85$ |
| **Middle: CS2 = Shock -> Reward** | | | |
| Sessions 1-8<br>Two-way mixed-effects model | Session<br>$F_{(1.90, 15.17)} = 3.11$, $p = 0.08$ | Trial<br>$F_{(1, 8)} = 1.82$, $p = 0.21$ | Interaction<br>$F_{(2.93, 21.75)} = 1.89$, $p = 0.16$ |
| Sessions 9-16<br>Two-way mixed-effects model | Session<br>$F_{(2.90, 23.18)} = 2.65$, $p = 0.07$ | Trial<br>$F_{(1, 8)} = 0.18$, $p = 0.68$ | Interaction<br>$F_{(2.67, 16.78)} = 0.33$, $p = 0.78$ |
| Sessions 17-24<br>Two-way mixed-effects model | Session<br>$F_{(2.14, 14.95)} = 2.89$, $p = 0.08$ | Trial<br>$F_{(1, 7)} = 0.37$, $p = 0.56$ | Interaction<br>$F_{(2.83, 12.56)} = 1.34$, $p = 0.30$ |
| **Right: CS-** | | | |
| Sessions 1-8<br>Two-way mixed-effects model | Session<br>$F_{(3.09, 24.74)} = 1.83$, $p = 0.17$ | Trial<br>$F_{(1, 8)} = 0.02$, $p = 0.87$ | Interaction<br>$F_{(3.27, 24.27)} = 1.53$, $p = 0.23$ |
| Sessions 9-16<br>Two-way mixed-effects model | Session<br>$F_{(3.35, 26.80)} = 0.48$, $p = 0.72$ | Trial<br>$F_{(1, 8)} = 0.40$, $p = 0.54$ | Interaction<br>$F_{(3.45, 21.66)} = 1.56$, $p = 0.22$ |
| Sessions 17-24<br>Two-way mixed-effects model | Session<br>$F_{(2.62, 18.33)} = 1.43$, $p = 0.27$ | Trial<br>$F_{(1, 7)} = 0.15$, $p = 0.71$ | Interaction<br>$F_{(2.70, 11.56)} = 1.88$, $p = 0.19$ |
| **Panel E: 5 trial bin CS response averaged across early and late phases of training** | | | |
| **Left: CS- early training phase** | | | |
| Dopamine<br>Two-way mixed-effects model | Training phase<br>$F_{(1.06, 7.42)} = 1.68$, $p = 0.24$ | Trials<br>$F_{(1, 7)} = 11.76$, ***p* = 0.01** | Interaction<br>$F_{(1.46, 10.24)} = 1.37$, $p = 0.29$ |
| GABA<br>Two-way mixed-effects model | Training phase<br>$F_{(1.30, 10.42)} = 0.70$, $p = 0.46$ | Trials<br>$F_{(1, 8)} = 4.01$, $p = 0.08$ | Interaction<br>$F_{(1.27, 8.86)} = 3.17$, $p = 0.10$ |
| **Right: CS- late training phase** | | | |
| Dopamine<br>Two-way mixed-effects model | Training phase<br>$F_{(1.59, 11.14)} = 2.01$, $p = 0.18$ | Trials<br>$F_{(1, 7)} = 3.39$, $p = 0.11$ | Interaction<br>$F_{(1.82, 12.73)} = 1.36$, $p = 0.29$ |
| GABA<br>Two-way mixed-effects model | Training phase<br>$F_{(1.31, 10.48)} = 0.30$, $p = 0.66$ | Trials<br>$F_{(1, 8)} = 8.36$, ***p* = 0.02** | Interaction<br>$F_{(1.70, 10.23)} = 1.83$, $p = 0.21$ |

| Table 4: Supplemental Figure 7 – mPFC-VTA chemogenetic inhibition during FCL | | | |
|---|---|---|---|
| **Panel A: Conditioned responding – 5 trial bin** | | | |
| **Left: $CS_1$ = Reward -> Shock** | | | |
| Sessions 1-8<br>Two-way mixed-effects model | Session<br>$F_{(2.17, 34.64)} = 11.42$, $\mathbf{p = 0.0001}$ | Trial<br>$F_{(1, 16)} = 18.67$, $\mathbf{p = 0.0005}$ | Interaction<br>$F_{(2.96, 47.01)} = 1.52$, $p = 0.22$ |
| Sessions 9-16<br>Two-way mixed-effects model | Session<br>$F_{(2.10, 33.60)} = 8.50$, $\mathbf{p = 0.0009}$ | Trial<br>$F_{(1, 16)} = 0.18$, $p = 0.68$ | Interaction<br>$F_{(3.87, 58.52)} = 3.65$, $\mathbf{p = 0.01}$ |
| Sessions 17-24<br>Two-way mixed-effects model | Session<br>$F_{(1.94, 27.09)} = 5.81$, $\mathbf{p = 0.009}$ | Trial<br>$F_{(1, 14)} = 7.98$, $\mathbf{p = 0.01}$ | Interaction<br>$F_{(4.17, 53.58)} = 2.70$, $\mathbf{p = 0.04}$ |
| **Middle: $CS_2$ = Shock -> Reward** | | | |
| Sessions 1-8<br>Two-way mixed-effects model | Session<br>$F_{(4.84, 77.43)} = 1.71$, $p = 0.14$ | Trial<br>$F_{(1, 16)} = 3.84$, $p = 0.07$ | Interaction<br>$F_{(4.10, 64.35)} = 0.83$, $p = 0.51$ |
| Sessions 9-16<br>Two-way mixed-effects model | Session<br>$F_{(2.08, 33.26)} = 8.53$, $\mathbf{p = 0.0009}$ | Trial<br>$F_{(1, 16)} = 28.73$, $\mathbf{p < 0.0001}$ | Interaction<br>$F_{(4.13, 62.56)} = 0.43$, $p = 0.79$ |
| Sessions 17-24<br>Two-way mixed-effects model | Session<br>$F_{(2.24, 31.30)} = 7.19$, $\mathbf{p = 0.002}$ | Trial<br>$F_{(1, 14)} = 0.92$, $p = 0.35$ | Interaction<br>$F_{(2.76, 35.54)} = 1.18$, $p = 0.33$ |
| **Right: CS-** | | | |
| Sessions 1-8<br>Two-way mixed-effects model | Session<br>$F_{(4.48, 71.67)} = 9.93$, $p = 0.46$ | Trial<br>$F_{(1, 16)} = 0.84$, $p = 0.37$ | Interaction<br>$F_{(4.84, 76.70)} = 1.08$, $p = 0.38$ |
| Sessions 9-16<br>Two-way mixed-effects model | Session<br>$F_{(3.81, 60.91)} = 0.41$, $p = 0.79$ | Trial<br>$F_{(1, 16)} = 2.28$, $p = 0.15$ | Interaction<br>$F_{(2.69, 40.76)} = 0.48$, $p = 0.68$ |
| Sessions 17-24<br>Two-way mixed-effects model | Session<br>$F_{(3.23, 45.21)} = 2.48$, $p = 0.07$ | Trial<br>$F_{(1, 14)} = 1.23$, $p = 0.29$ | Interaction<br>$F_{(3.35, 43.06)} = 0.33$, $p = 0.82$ |
| **Panel B: 5 trial bin conditioned responding averaged across early and late phases of training** | | | |
| **Left: Early training phase** | | | |
| $CS_1$ = Reward -> Shock<br>Two-way mixed-effects model | Training phase<br>$F_{(1.27, 20.33)} = 7.09$, $\mathbf{p = 0.01}$ | Trials<br>$F_{(1, 16)} = 10.58$, $\mathbf{p = 0.005}$ | Interaction<br>$F_{(1.85, 25.91)} = 8.96$, $\mathbf{p = 0.001}$ |
| $CS_2$ = Shock -> Reward<br>Two-way mixed-effects model | Training phase<br>$F_{(1.38, 22.08)} = 16.74$, $\mathbf{p = 0.0002}$ | Trials<br>$F_{(1, 16)} = 4.21$, $p = 0.06$ | Interaction<br>$F_{(1.67, 23.32)} = 5.65$, $\mathbf{p = 0.01}$ |
| CS-<br>Two-way mixed-effects model | Training phase<br>$F_{(1.28, 20.46)} = 2.49$, $p = 0.12$ | Trials<br>$F_{(1, 16)} = 2.74$, $p = 0.12$ | Interaction<br>$F_{(1.70, 23.84)} = 0.50$, $p = 0.58$ |
| **Right: Late training phase** | | | |
| $CS_1$ = Reward -> Shock<br>Two-way mixed-effects model | Training phase<br>$F_{(1.90, 30.47)} = 14.13$, $\mathbf{p < 0.0001}$ | Trials<br>$F_{(1, 16)} = 5.53$, $\mathbf{p = 0.03}$ | Interaction<br>$F_{(1.53, 21.39)} = 1.30$, $p = 0.28$ |
| $CS_2$ = Shock -> Reward<br>Two-way mixed-effects model | Training phase<br>$F_{(1.05, 16.81)} = 14.65$, $\mathbf{p = 0.001}$ | Trials<br>$F_{(1, 16)} = 6.88$, $\mathbf{p = 0.02}$ | Interaction<br>$F_{(1.61, 22.48)} = 10.91$, $\mathbf{p = 0.0009}$ |
| CS-<br>Two-way mixed-effects model | Training phase<br>$F_{(1.61, 25.75)} = 3.15$, $p = 0.07$ | Trials<br>$F_{(1, 16)} = 1.65$, $p = 0.22$ | Interaction<br>$F_{(1.81, 25.39)} = 0.74$, $p = 0.47$ |
